# The earliest circadian clock in the mammalian brain emerges in the embryonic choroid plexus

**DOI:** 10.1101/2025.09.09.675224

**Authors:** Hélène Vitet, Vuong Hung Truong, Jihwan Myung

## Abstract

Maternal chronodisruption is linked to miscarriages and neurodevelopmental risks, yet the earliest brain circadian oscillator in the embryo and the mechanism of its emergence remain unknown. Using bioluminescent reporters for PER2 and *Bmal1*, we tracked the onset of autonomous ∼24 h oscillations *ex vivo*. While the suprachiasmatic nucleus (SCN) is widely considered the first circadian structure at embryonic day (E) 14.5-15.5, we find that the fourth ventricle choroid plexus (4VCP) exhibits PER2 oscillations by E11.5-E12.5. These oscillations show hallmarks of a saddle-node on an invariant circle (SNIC) bifurcation, characterized by an initially long period converging toward 24 h and pronounced sensitivity to sub-degree temperature cycles mimicking physiological maternal cues. Transcriptional profiling of maternal and embryonic 4VCPs unveiled three epochs in clock gene expression that parallel tissue differentiation. We identify the 4VCP as the earliest detectable brain circadian oscillator, and demonstrate how its bifurcation dynamics allow the nascent clock to couple to maternal rhythms.

## Introduction

Circadian clocks are autonomous pacemakers that align physiological processes with Earth’s 24-h cycle, maintained by a robust transcription-translation feedback loop (TTFL). The transcription factors BMAL1 and CLOCK heterodimerize to activate the transcription of negative feedback components, including the *Period* (*Per1/2/3)* and *Cryptochrome* (*Cry1/2*) genes. The resulting PER and CRY proteins accumulate, form complexes, and undergo post-translational modifications that introduce a delay that sets the circadian timescale. These complexes translocate into the nucleus to repress their own transcription to complete the loop. The transcription of *Bmal1* is further regulated by REV-ERB and ROR, conferring additional robustness to the oscillation. The assemblage of these loops creates tissue-specific phase organization of clock gene expression (Myung et al., 2025). In adult mammals, the suprachiasmatic nucleus (SCN) in the hypothalamus acts as the central clock, synchronizing peripheral clocks throughout the body (Finger et al., 2020). Understanding when and how this system first emerges during development is important, as maternal chronodisruption is associated with preterm birth, miscarriage, and adverse neurodevelopmental outcomes (Astiz and Oster, 2021); yet, neither the identity nor the onset mechanism of the earliest embryonic brain clock has been established.

Circadian rhythms have been detected prenatally in humans, non-human primates, and rodents (Serón-Ferré et al., 2012; Landgraf et al., 2014). However, detection is technically challenging, due to low-amplitude oscillations and tissue-specific differences in clock gene expression. Although *Per* and *Bmal1* show no clear rhythms in whole-embryo preparations (E10-E17) (Cao et al., 2022; Dolatshad et al., 2010; Serón-Ferré et al., 2012), circadian rhythmicity of PER2 has been detected in the SCN *ex vivo* around E14.5 to E15.5 (Wreschnig et al., 2014; Carmona-Alcocer et al., 2018). Whether the SCN is truly the first circadian structure in the embryonic brain, however, has not been systematically examined.

A strong candidate for an even earlier clock is the choroid plexus (CP). In adult mice, the CP acts as a robust clock capable of entraining the SCN through a diffusive pathway (Myung et al., 2018). This non-neuronal tissue lines the brain ventricles and filters blood to produce cerebrospinal fluid (CSF) and a distinct secretome (Saunders et al., 2023; Chau et al., 2015; Courtney et al., 2025; Edelbo et al., 2023). Among the CPs, the fourth ventricle CP (4VCP) is the first to develop, arising from the lower rhombic lip as early as E9.5, expressing *Wnt1*, *Ttr*, and *Kcne2*, with development rate peaking at E11-E12.5 (Lun et al., 2015; Hunter and Dymecki, 2007). By E14, the 4VCP becomes anatomically apparent and is thought to be differentiated, although proliferation continues postnatally (Lun et al., 2015).

Beyond mechanical support, CSF is a key volumetric regulator of early brain development. Embryonic CSF is present at neural tube closure (E9.5), initially derived from enclosed amniotic fluid before transitioning to CP-produced fluid between E10.5 and E14.5 (Chau et al., 2015). Because CP-derived volume transmission dominates early brain-wide signaling (E10.5-E11.5) and the adult CP secretome is under circadian control (Edelbo et al., 2023), an embryonic CP clock could provide early time cues for neural progenitor programs (Miyan et al., 2003; Martin et al., 2009; Gato et al., 2005).

How might such a clock emerge? The early embryo operates on timescales of hours for cell division and somitogenesis, far shorter than the 24 h cycle, and an abrupt transition from the “somitogenesis era” to the “circadian era” has been proposed (Caviness et al., 1995; Yagita, 2024). From a nonlinear dynamics perspective, a quiescent state can transition to stable oscillation through a bifurcation. Circadian clocks, along with many oscillations in biology, are often modelled as a supercritical Hopf bifurcation, in which amplitude grows gradually while the period remains constant (Murayama et al., 2017). However, developmental transitions, such as the onset of the first heartbeat in zebrafish, can arise via a noisy saddle-node on an invariant circle (SNIC) bifurcation (Jia et al., 2023). A SNIC produces finite amplitude oscillations abruptly, with an initially long period that shortens toward ∼24 h. Near threshold, the slow dynamics makes oscillation frequency highly sensitive to weak external forcing, so that even small maternal cues could effectively entrain a nascent clock. Distinguishing between these two mechanisms requires tracking both period and amplitude across development, which have not previously been available for any embryonic circadian tissue.

In this study, we pinpoint when circadian rhythms first emerge in the embryonic mouse brain. Using bioluminescent reporters for PER2 and *Bmal1*, we identify that the 4VCP is the earliest detectable clock, governed by SNIC-like bifurcation dynamics. We demonstrate that this onset mechanism renders the nascent clock highly sensitive to mid-gestational fluctuations in the maternal clock. Transcriptional profiling of mother and embryo throughout pregnancy reveals three distinct epochs in clock gene expression coinciding with 4VCP differentiation. Initially coupled to the mother, both maternal and embryonic clocks undergo transient destabilization, after which the embryo establishes autonomous circadian organization approaching canonical TTFL architecture. As the 4VCP matures, it ceases to entrain to external temperature cycles and transitions toward robust, self-sustained oscillation.

## Results

### The 4VCP exhibits autonomous circadian oscillations prior to the SCN

The embryonic SCN is widely recognized as the earliest reported brain region to exhibit a detectable circadian rhythm, emerging between E14.5 and E15.5 (Wreschnig et al., 2014; Carmona-Alcocer et al., 2018). To test whether this holds across the brain, we prepared brain slices from E15.5 PER2::LUC^+/-^ embryos and performed bioluminescence imaging for 5-7 days (**Fig. 1A**-**C**, **Movie S1**). Spectral analysis revealed two consistent loci with ∼24 h periods, circadian power exceeding 30% (see Materials and Methods), and high RMS amplitude (**Fig. 1Ca,1Cb**, **Fig. S4**). The ventral locus (white arrow) was RORA-positive, confirming its identity as the SCN (VanDunk et al., 2011), whereas the posterior structure (red arrow) was TTR-positive, a known marker for the CP (Kaiser and Bryja, 2020) (**Fig. 1Cc**). When analysis was restricted strictly to these regions, robust rhythms with periods between 20 and 28 h were detected, with circadian power of 59.4 ± 0.8% in the SCN and 40.4 ± 0.4% in the 4VCP (**Fig. 1Cd**). The pituitary gland also exhibited rhythmicity (46.9 ± 0.4%), but with lower RMS amplitude (10.2 ± 0.2 AU vs. SCN: 15.5 ± 0.9 AU, 4VCP: 15.5 ± 0.5 AU; both *p* < 0.0001) (**Fig. S4**).

**Figure 1.**
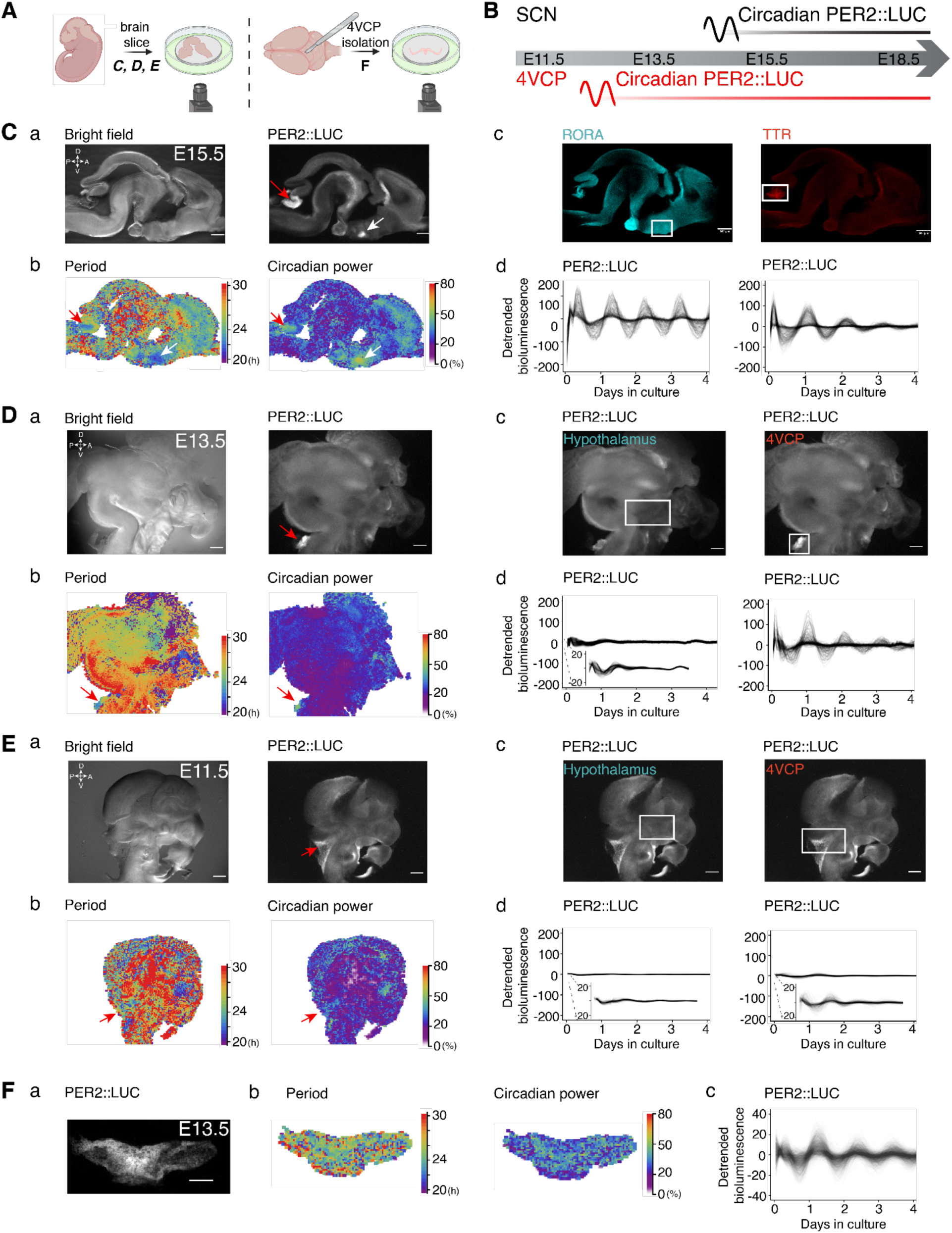
Location of the first clock in the mouse embryonic brain. (**A**) Workflow of PER2::LUC bioluminescence recording from embryonic brain/head slices (left; C-E) or isolated 4VCP explants (right; F). (**B**) Developmental timeline summarizing that 4VCP PER2::LUC rhythms precede those in the SCN. (**C**) E15.5 PER2::LUC brain slice. **a**: Bright-field (left) and bioluminescence (right) images. **b**: Maps of FFT-derived period (left) and circadian power (right; see Materials and Methods). **c**: Post hoc immunofluorescence for RORA (left) and the CP marker TTR (right) after tissue clearing. **d**: Detrended PER2::LUC time series from the regions indicated by white boxes in Cc; see **Movie S1**. Representative of 3 experiments (5 slices total). (**D**) E13.5 PER2::LUC head slice. **a**: Bright-field (left) and bioluminescence (right) images. **b**: Period (left) and circadian power (right) maps as in Cb. **c**: White boxes indicate the putative hypothalamus/SCN region (left) and 4VCP (right). **d**: Detrended time series from the regions in Dc; see **Movie S2**. Representative of 3 experiments (4 slices total). (**E**) E11.5 PER2::LUC head slice. **a**: Bright-field (left) and bioluminescence (right) images. **b**: Period (left) and circadian power (right) maps. **c**: White boxes indicate putative hypothalamus (left) and 4VCP (right). **d**: Detrended time series from the regions in Ec; see **Movie S3**. Representative of 3 experiments (3 slices total). (**F**) Isolated E13.5 PER2::LUC 4VCP explant. **a**: Bioluminescence image. **b**: Period (left) and circadian power (right) maps. **c**: Detrended time series over 4 days; see **Movie S4**. Representative of 4 experiments (7 explants total). Throughout, red arrows indicate the 4VCP; white arrows indicate the SCN. Scale bar: 500 µm.

At E13.5, no circadian oscillations were detected in ventral structures corresponding to SCN progenitors, consistent with previous reports (Wreschnig et al., 2014; Carmona-Alcocer et al., 2018) (**Fig. 1D**, **Movie S2**). However, the 4VCP (red arrow) exhibited a near-24 h rhythm with strong circadian power of 30.0 ± 0.7% (vs. hypothalamus: 20.7 ± 0.1%) and high RMS amplitude (**Fig. 1Db**, **Fig. S4**), which we confirmed after restricting the analysis to the 4VCP (**Fig. 1Dd**). At E11.5, no significant circadian rhythm was detected in whole brain slices (**Fig. 1E**, **Movie S3**). To confirm that this rhythm was endogenous, we dissected E13.5 4VCPs and recorded bioluminescence *ex vivo* (**Fig. 1F**, **Movie S4**). The isolated 4VCP explant exhibited robust rhythmicity with circadian power of ∼40%, well above the detection threshold determined from blank luciferase medium (Blk, < 20%) (**Fig. 1Fb**).

### Tracking the precise timeline of 4VCP clock emergence

To pinpoint the exact onset of clock emergence, we tracked PER2::LUC bioluminescence in isolated 4VCPs from E9.5, when the CSF compartment first forms (Chau et al., 2015), through to postnatal stages via LumiCycle, recording from 110 embryos (3-11 per stage, from at least 2-3 litters) for 5-7 days (**Fig. 2**). The oscillation period was initially long and highly dispersed at E9.5 (27.50 ± 2.12 h; *p* < 0.0001, *n* = 3 detectable oscillations), converging toward a stable 24 h by E11.5 (24.98 ± 0.54 h; *p* > 0.1, *n* = 11). This marked the onset of detectable oscillations, as circadian power exceeded 20% (31.42 ± 1.90% at E11.5, *p* < 0.0001, *n* = 15; vs. 18.01 ± 2.43% at E10.5, *p* > 0.05, *n* = 15) (**Fig. 2A**,**C****,D**). These early oscillations were weakly sustained with low RMS amplitude. By E12.5, they persisted throughout recording, with amplitude increasing further at E13.5. The rhythm became fully circadian at E15.5 (24.47 ± 0.60 h; *n* = 5), with high circadian power (50.96 ± 3.42%; *n* = 5) approaching the 2-day window ceiling of ∼60% (**Fig. 2A,C,D**).

**Figure 2.**
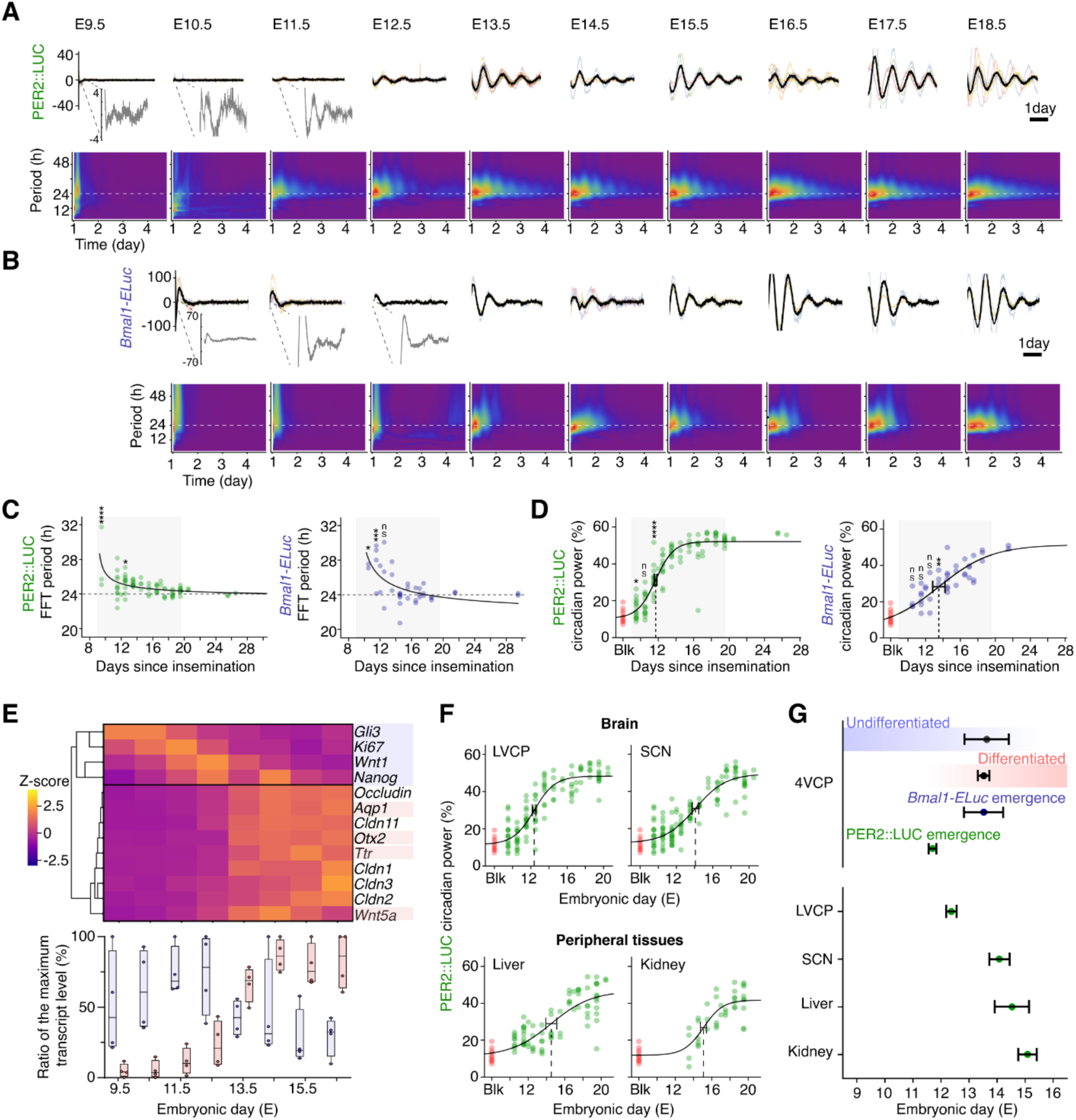
Quantification of clock emergence in brain and peripheral tissues, and characterization of 4VCP differentiation. (**A**) Individual (thin colored) and mean (black) detrended PER2::LUC bioluminescence traces (top) and spectrograms (bottom) from the embryonic 4VCP, E9.5 to E18.5. Zoom-in insets highlight low-amplitude rhythms at early stages. (**B**) As in (A), for *Bmal1-ELuc*, from E10.5 to E18.5. (**C**) Period scatter plots for PER2::LUC (left) and *Bmal1-ELuc* (right) with 1/√(*t* − *t*₀) fits. Asterisks indicate significant differences from 24 h (ns, not significant; \**p* < 0.05; \*\*\**p* < 0.001; \*\*\*\**p* < 0.0001). (**D**) Circadian power scatter plots for PER2::LUC (left) and *Bmal1-ELuc* (right) with sigmoidal fits; emergence times (*T_e_*) were E11.7 ± 0.1 and E13.5 ± 0.7, respectively (best fit estimate ± SE). Asterisks indicate significant differences from 20% (as in (C)). (**E**) Heatmap of developmental marker genes (top) and box-and-whisker plots (bottom) comparing stem/progenitor (blue) and differentiation (red) markers across embryonic stages; each gene is normalized to its maximum expression between E9.5 and E16.5. Each data point represents the mean of 6-16 litters (3-11 embryos per litter). Boxes show interquartile range (25th and 75th percentile) and median; whiskers indicate minimum and maximum values. (**F**) Developmental trajectories of PER2::LUC circadian power in the LVCP and the SCN (top) and liver and kidney (bottom), with sigmoidal fits. (**G**) Comparison of half-maximal times for progenitor markers (E13.6 ± 0.8) and differentiation markers (E13.5 ± 0.2) with emergence times of *Bmal1-ELuc* and PER2::LUC oscillations in the 4VCP, and PER2::LUC emergence time for each tissue. Error bars: mean ± SEM after outlier exclusion.

To characterize the molecular assembly of this clock, we also measured from the *Bmal1-ELuc* reporter strain. Interestingly, *Bmal1-ELuc* oscillations with a ∼24 h period were only observed from E13.5 to E14.5, approximately two days after the emergence of PER2 oscillations (**Fig. 2B,C,D**). Sigmoidal fits to circadian power trajectories estimated the exact emergence time (*T_e_*) (see Materials and Methods). PER2::LUC oscillations became circadian at E11.7 ± 0.1, whereas *Bmal1-ELuc* oscillations emerged later at E13.5 ± 0.7 *ex vivo* (**Fig. 2D**). This delayed *Bmal1* aligns with the sudden upward shift in PER2::LUC circadian power above 20% observed at E12.5-E13.5 (**Fig. S5A**).

Because the circadian clock and developmental programs are deeply intertwined, we mapped the embryonic 4VCP’s developmental trajectory by quantifying progenitor markers (*Gli3*, *Ki67*, *Wnt1*, *Nanog*) (Awatramani et al., 2003; Hunter and Dymecki, 2007) and differentiation markers (*Aqp1*, *Otx2, Ttr*, *Wnt5a*) (Lun et al., 2015; Langford et al., 2020). Progenitor markers predominated from E9.5 to E11.5, whereas differentiation genes dominated from E14.5 onward (**Fig. 2E**). Sigmoidal fits estimated the progenitor half-minimum at E13.6 ± 0.8 and the differentiation half-maximum at E13.5 ± 0.2, coinciding with *Bmal1-ELuc* emergence (*T_e_* = E13.5 ± 0.7) (**Fig. 2G**). This defines three epochs: the undifferentiated state (E9.5-E12), the transition state (E12-E15), and the differentiated state (E15-E16.5). Notably, PER2::LUC oscillations (*T_e_* = E11.7 ± 0.1) appear during the early transition, preceding both *Bmal1* rhythmicity and the differentiation switch, an order that is atypical for a canonical TTFL where *Bmal1* drives *Per* transcription (**Fig. 2G**).

Finally, we compared the emergence time of PER2::LUC rhythms across the brain and peripheral organs in isolated explant culture. Among all tissues tested, the 4VCP was the first to become circadian, preceding the lateral ventricle CP (LVCP: E12.4 ± 0.2), the SCN (E14.2 ± 0.4), and peripheral organs such as the liver (E14.5 ± 0.6) and kidney (E15.1 ± 0.3) (**Fig. 2F,G**). In the SCN, the period approached 24 h at E14.5 (24.47 ± 2.35 h, *n* = 7) and became fully circadian at E15.5 with high circadian power (44.56 ± 2.48%, *n* = 3).

### Abrupt, SNIC-like onset and synchronization of clock emergence

The type of bifurcation underlying oscillation onset can be deduced by tracking how amplitude and period evolve over development. Circadian clocks have long been modelled as supercritical Hopf bifurcations, which behave analogously to resonance: amplitude grows smoothly while the period remains fixed. A SNIC bifurcation is fundamentally different and oscillations appear abruptly at full amplitude but with an initially long period that shortens toward ∼24 h. SNIC oscillation also displays pronounced phase sensitivity to weak perturbations (Ermentrout, 1996). In single-cell PER2::LUC imaging, we observed a sharply defined clock onset with a stepwise amplitude increase (**Fig. 3A**) and an initially long period (**Fig. 3B**), hallmarks of a SNIC bifurcation. Information-theoretic model comparison favored a discrete amplitude step and period decay, consistent with SNIC dynamics rather than the gradual emergence expected from a Hopf bifurcation (**Fig. S5C**). This was further supported at the tissue level: whole-tissue luminometry confirmed a long initial period at E9.5 (26.4 ± 4.3 h), which converged to ∼24 h by E17.5 (24.3 ± 1.1 h) (**Fig. 2C**), and period scatter plots fit the SNIC scaling law 1/√(*t* − *t*₀). While the whole-tissue luminometry amplitude increased linearly, this reflected growth in 4VCP cell number across development (**Fig. S5B**). *In vivo*, *Per2* transcript periods similarly reached ∼28 h at E9.5 before converging toward 24 h (**Fig. S8**).

**Figure 3.**
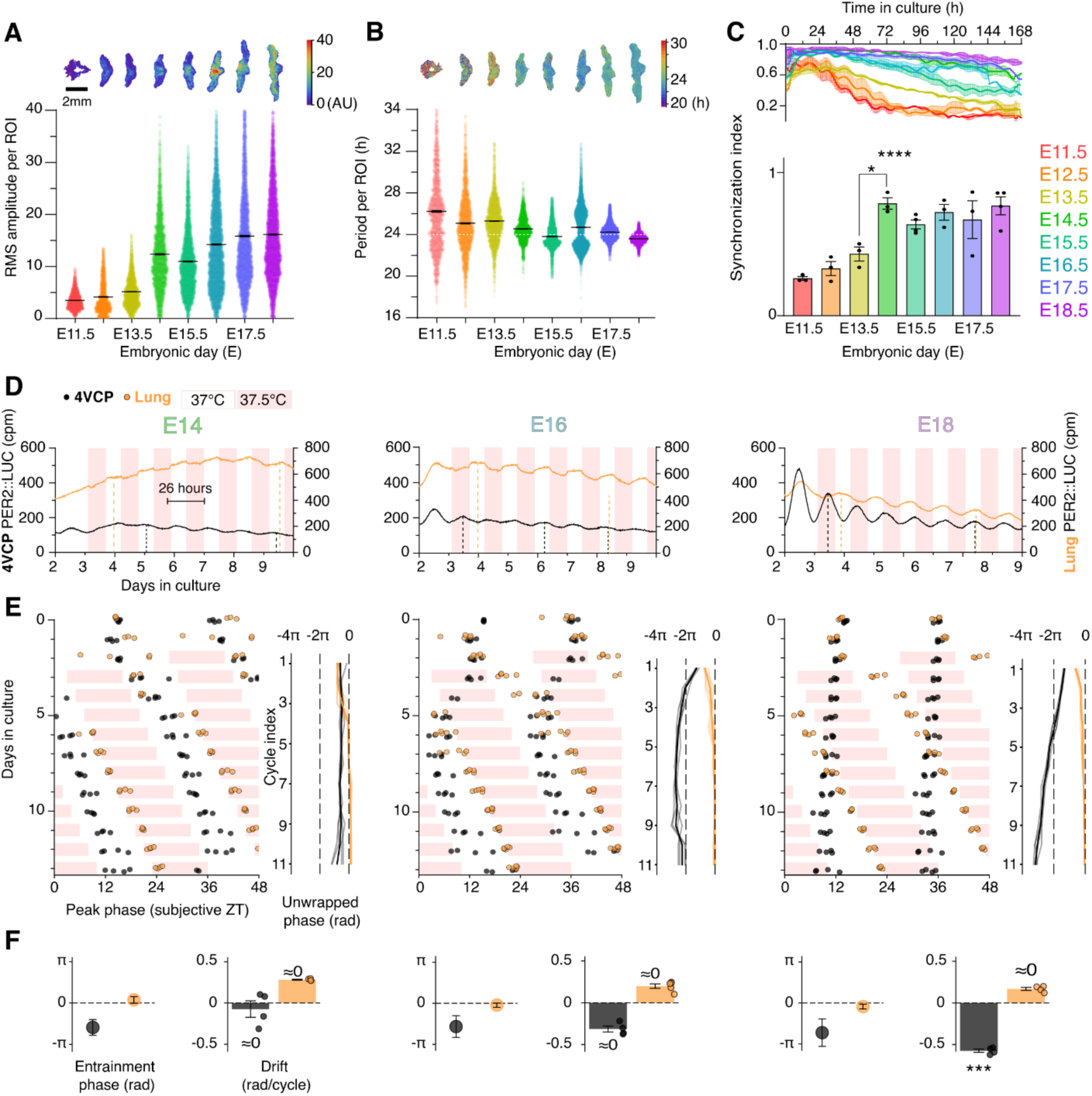
Characterization of the bifurcation type describing the onset of the embryonic 4VCP clock. (**A**) Single-cell RMS amplitude and (**B**) period of PER2::LUC in the embryonic 4VCP from E11.5 to E18.5. Insets above each panel show representative spatial maps across stages (scale bar: 2 mm). *n* = 579-2,787 ROIs from 3-4 4VCP explants and 2-3 litters per stage, after outlier exclusion. White dotted line indicates 24 h. (**C**) Synchronization index (Kuramoto order parameter) in the embryonic 4VCP explant, as it evolves over days in culture (top) and time-averaged individual values across stages, overlaid with scatter plots (bottom; same *n* as in (A)). One-way ANOVA (*F*(7,18) = 10.72, *p* < 0.0001) with Tukey’s post hoc test (\**p* = 0.0194, \*\*\*\**p* < 0.0001). (**D**) Representative raw PER2::LUC time series from the 4VCP (black) and lung (orange) at E14 (left), E16 (middle), and E18 (right). Pale red shading indicates 37.5 °C epochs. (**E**) Double-plots of PER2::LUC peak phase from the corresponding explants, alongside unwrapped phase plots (rad). (**F**) Entrainment phase (rad) and phase drift (rad/cycle) for the 4VCP and lung across the three developmental stages. Phase drift was assessed using equivalence testing (TOST; bounds ±0.483 rad/cycle) together with one-sample *t*-tests (see Methods). At E18, 4VCP displayed significant negative drift (-0.473 ± 0.042 rad/cycle, *p* = 0.0015), whereas the lung remained equivalent to zero. Error bars: mean ± SEM after outlier exclusion.

At the population level, we observed an abrupt rise in cellular synchronization between E13.5 and E14.5 (**Fig. 3C**, **Fig. S6A**) quantified by the Kuramoto order parameter (0 indicates the absence of synchronization; 1 denotes complete synchronization). This shift parallels the sudden synchronization onset reported in the SCN (Carmona-Alcocer et al., 2018). These changes demonstrate that the clock emergence process at both single-cell and population levels is not gradual ramp-up, but rather a discretely staged event occurring in specific developmental epochs.

### Temperature cycles drive embryonic 4VCP rhythmicity until E16.5

Since systems near a SNIC bifurcation are highly sensitive to low-amplitude stimuli, we hypothesized that subtle maternal circadian temperature cycles could serve as an early, non-blood-borne driver of the nascent clock. During pregnancy, the amplitude of the maternal core body temperature rhythm dampens to ∼1 °C and further drops to ∼0.5 °C by mid-gestation (Smarr et al., 2016). This ∼0.5 °C fluctuation is considered too small to entrain adult peripheral clocks *in vitro* (∼4 °C, Brown et al., 2002; ∼2.5 °C, Buhr et al., 2010; ∼1.5 °C, Abraham et al., 2010).

To test this directly, we recorded PER2::LUC activity in cultured embryonic explants subjected to temperature entrainment. Since maternal body temperature at E6 varies between 37.5 °C and 37 °C (Wharfe et al., 2016), we applied an initial temperature of 37 °C for 2 days, followed by 12 days of a 37-37.5 °C (Δ0.5 °C) square-wave cycle with a 26 h period (**Fig. 3D**). We evaluated explants of the 4VCP and lung at E14, E16, and E18 and observed a clear, stage-dependent transition in entrainability (**Fig. 3E**). At E14, the 4VCP PER2 rhythm phase-locked to the 26 h temperature cycle (circular SD = 0.47 rad; *n* = 5). At E16, entrainment occurred only after an initial transient (circular SD = 0.70 rad; *n* = 4). By E18, the 4VCP was resistant to entrainment, failing to phase-lock to the 26 h cycle (circular SD = 1.16 rad; *n* = 4) (**Fig. 3F**). The lung entrained at all stages, reflecting its prolonged maturation and weaker intercellular coupling (Abraham et al., 2010) (**Fig. 3E,F**). The LVCP mirrored the 4VCP, losing entrainability by E18, whereas the SCN remained entrainable at E18, consistent with its network immaturity at this stage (Bedont and Blackshaw, 2015) (**Fig. S6B**, **Table S2**).

By E18, the 4VCP behaved as a robust, adult-like limit cycle oscillator, resistant to small perturbations. Despite receiving higher-amplitude blood-borne signals from ∼E16 onward, the fully differentiated 4VCP effectively “closed” its window of extreme sensitivity to low-amplitude ambient fluctuations to operate as an independent pacemaker.

### Transcriptional profiling reveals non-canonical early rhythms in three epochs

We sampled embryonic 4VCPs from 127 PER2::LUC litters across E9-E16 at four circadian time points (ZT2, ZT8, ZT14, ZT20) per embryonic day for RT-qPCR. The highest transcript levels were those of *Clock, Rora,* and *Chrono*, rather than *Bmal1* and *Per1/2/3* as expected from adult TTFL networks (**Fig. 4A**). Transcripts followed distinct developmental trajectories: some increased gradually (*Clock, Dbp, Rev-Erbα/β, Bmal1*, *Per1*), others peaked sharply around E14.5 (*Rora, Chrono*, *Per2/3*), and a few remained relatively stable (*Rorc*, *Cry2*). The divergent trajectories of *Bmal1* and *Per2* likely explain the onset-timing discrepancy between *Bmal1-ELuc* and PER2::LUC reporters (**Fig. 2G**).

**Figure 4.**
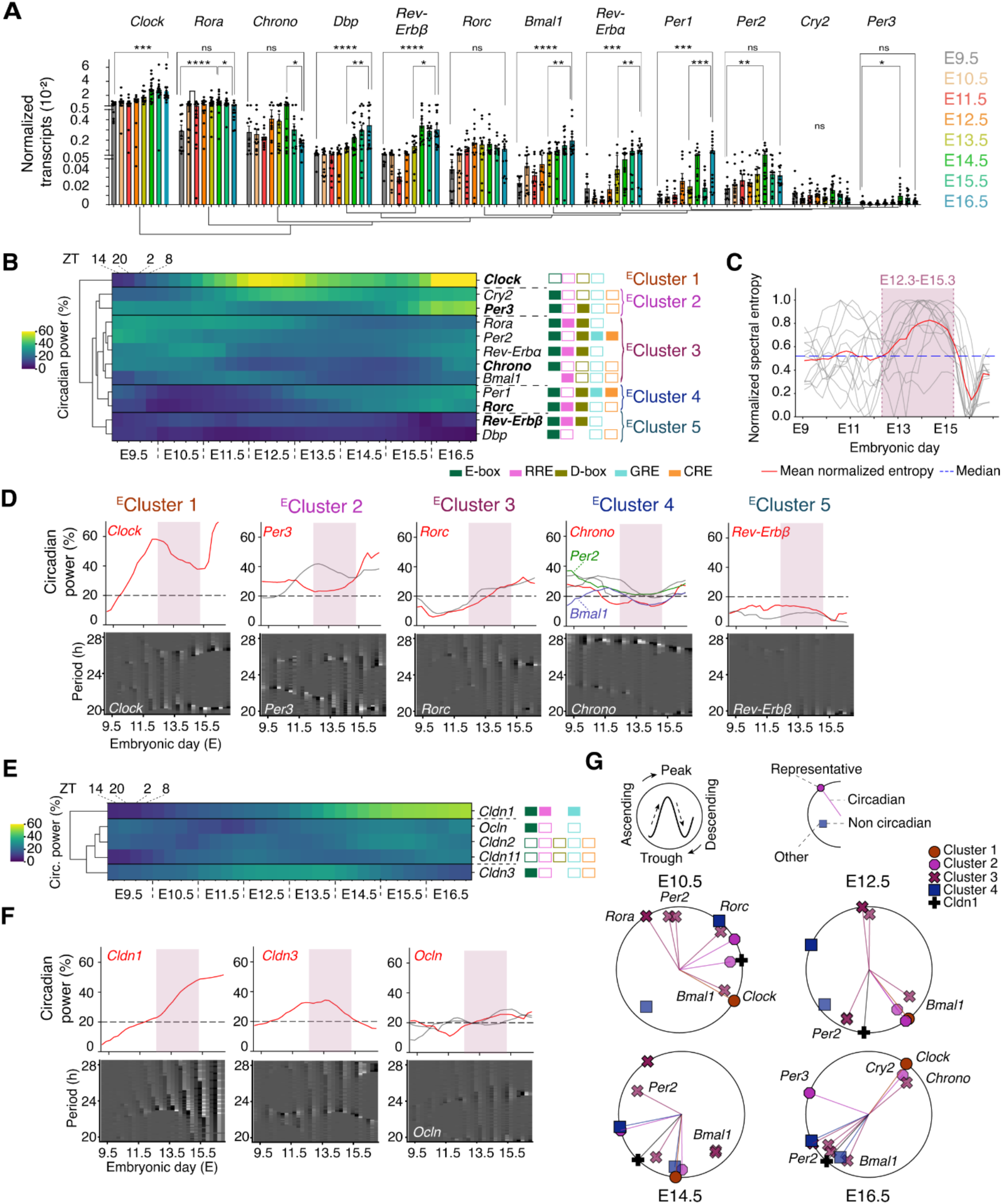
Rhythmic expression of clock genes in the embryonic 4VCP across development. (**A**) Normalized mRNA abundance of core clock genes in the embryonic 4VCP from E9 to E16 (*n* = 8-25 litters per stage). One-way ANOVA followed by Tukey’s post hoc test (ns, not significant; \**p* < 0.05; \*\**p* < 0.01; \*\*\**p* < 0.001; \*\*\*\**p* < 0.0001). (**B**) Hierarchical clustering and heatmap of embryonic circadian power of clock genes across stages and ZT. Tiles on the right annotate promoter-motif classes: filled tiles indicate validated motifs; outlined tiles indicate predictions (see **Fig. S7B**). Each data point represents the mean of 2-5 litters (3-11 embryos per litter). (**C**) Normalized spectral entropy across embryonic days. The mean entropy (red line) is plotted against the overall median (blue dashed line). The shaded region marks significant high-entropy epochs identified by permutation clustering: contiguous periods above the median that exceed the 95th percentile of cluster sizes from 1,000 random permutations (E12.3-E15.3). (**D**) Circadian power trajectories for each clock gene, grouped by cluster (top; representative gene in red; dashed line, 20% detection threshold). Shaded overlays indicate the high-entropy epoch (E12.3-E15.3). Circadian phase-weighted power spectrograms for the representative gene of each cluster across embryonic days (bottom). (**E**) As in (B), for tight junction genes (see also **Fig. S9B**). Each data point represents the mean of 1-4 litters. (**F**) As in (D), for tight junction genes. (**G**) Circular phase plots of embryonic gene expression across developmental stages. Peak phase is at 12 o’clock; phase rotates clockwise. Genes with circadian power ≥ 20% are connected to the center by lines and shown as opaque markers on the circle; non-circadian genes (< 20%) are shown as filled squares. Marker shape and color indicate cluster membership; the representative gene of each cluster is positioned on the outer radius. “^E^Cluster” denotes clusters derived from embryonic tissues. Error bars: mean ± SEM after outlier exclusion.

To characterize developmental evolution of the molecular rhythms, we quantified the circadian power of each transcript using SST-CWT analysis (Materials and Methods; **Fig. S7A**, **S8**). Hierarchical clustering of circadian power revealed five clusters with unique onset patterns (**Fig. 4B**): *Clock* (Cluster 1) becomes highly circadian from E9.5; *Per3*/*Cry2* (Cluster 2) are circadian early on (E9.5-E10.5) with a maximum power of ∼40-50%; *Rorc*/*Per1* (Cluster 3) emerge as circadian later, around E13.5; *Rora*/*Rev-Erbα*/*Per2*/*Chrono*/*Bmal1* (Cluster 4) show early rhythmicity from E9.5 (except for *Bmal1*) but lose it during the E12-E15 epoch; *Dbp*/*Rev-Erbβ* (Cluster 5) lack circadian rhythmicity across all measured stages. Several clock genes thus oscillate around E10.5, well before structural differentiation. Surprisingly, these empirical clusters do not align with the promoter-motif architecture expected from canonical TTFL logic (Ukai-Tadenuma et al., 2008), nor with the normalized density of transcription factor binding motifs quantified near their transcription start site (**Fig. S7B**). For instance, *Per1* and *Per2* both contain glucocorticoid and cAMP response elements (GREs and CREs) that can reset the CP clock (Liška et al., 2021), yet they fall into completely different onset clusters. This suggests that early embryonic circadian gene expression dynamics in the 4VCP cannot be solely explained by the transcriptional logic of the TTFL.

Spectral analysis across stages revealed unstable periods, visible as broad smears for most genes between E12 and E15 (**Fig. 4C**, **Fig. S7A,S8**), contrasting with the stable *Bmal1* period in maternal 4VCP (**Fig. 5B**, **Fig. S10A**). To quantify this globally, we calculated normalized spectral entropy, where high values indicate power dispersed across frequencies. Mean entropy peaked between E12.3 and E15.3, confirming a significant transition epoch (permutation clustering, *p* < 0.05; **Fig. 4C**).

**Figure 5.**
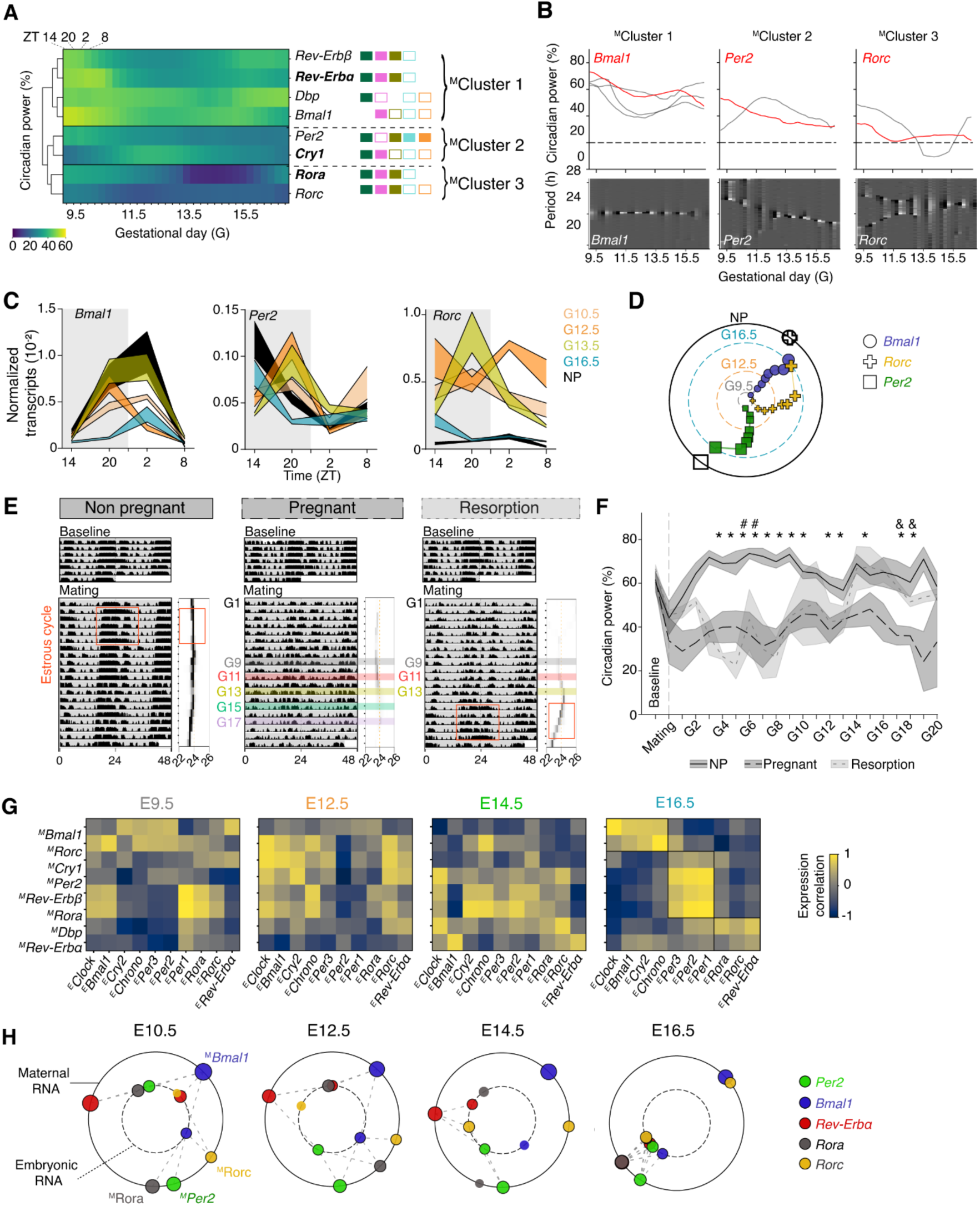
Maternal clock and maternal-embryonic transcriptional coordination across development. (**A**) Hierarchical clustering and heatmap of circadian power of maternal clock genes across gestational days and ZT. Tiles on the right annotate promoter-motif classes: filled tiles indicate validated motifs; outlined tiles indicate predictions (see **Fig. S7B**). Each data point represents the mean of 2-4 dams. (**B**) Circadian power trajectories for each maternal clock gene, grouped by cluster (top; representative gene in red; dashed line, 20% detection threshold). Circadian phase-weighted power spectrogram for *Bmal1, Per2* and *Rorc* across gestational days (bottom). (**C**) Normalized mRNA abundance of maternal *Bmal1*, *Per2* and *Rorc* across ZT at gestational days G10.5, G12.5, G13.5, and G16.5 (*n* = 2-4 dams) and in non-pregnant (NP) females (*n* = 3-5; black). Full time courses from G9.5 to G16.5 are shown in **Fig. S10B**. (**D**) Phase plot of maternal *Bmal1* (circle), *Per2* (square), and *Rorc* (cross) mRNA expression compared with non-pregnant female mice. Inner ring: G9.5 (grey), G12.5 (orange), and G16.5 (blue); outer ring: non-pregnant mice (NP). (**E**) Representative double-plotted actograms of a non-pregnant mouse, a pregnant mouse, and a pregnant mouse undergoing spontaneous resorption in DD (top: baseline before mating; bottom: after mating), with period and spectral power indicated by shading intensity (right panels; darker = higher power). (**F**) Circadian power of maternal locomotor activity throughout pregnancy (*n* = 5), spontaneous resorption (*n* = 2), and non-pregnant controls (*n* = 5). Two-sample *t*-test: *pregnant vs. non-pregnant (-5.4 < *t* < -2.8; 0.001 < *p* < 0.035); ^#^non-pregnant vs. resorption (*t* = 11.4, *p* = 0.0001 at G4; *t* = 4.3, *p* = 0.047 at G5), ^&^pregnant vs. resorption (*t* = -3.7; *p* = 0.014 at G17; *t* = -2.7, *p* = 0.043 at G18). (**G**) Heatmaps of pairwise correlation between maternal and embryonic clock genes at E9.5, E12.5, E14.5, and E16.5. (**H**) Dual-radius phase plots of maternal and embryonic gene expression across stages. Inner ring: embryonic cluster representatives; outer ring: selected maternal genes. Dotted lines connect maternal-embryonic pairs when both exhibit circadian power ≥ 20% and their phase difference is < *π*/2 rad (≤ 6 h). “^M^Cluster” denotes a cluster derived from maternal tissue. Error bars: mean ± SEM after outlier exclusion.

To investigate the functional consequences of this early clock, we tracked the expression of clock-controlled genes (CCGs) encoding tight junctions (*Cldn1/2/3/11*, *Ocln*) known to be present in both adult and embryonic CP epithelium (Wolburg et al., 2001; Steinemann et al., 2016). Functional E-boxes have been identified in several tight junction genes (Mei et al., 2015; Oh-oka et al., 2014), and we confirmed the robust expression of these transcripts in the embryonic 4VCP from E9.5 onwards (**Fig. S9A**). Consistent with this, *Cldn1* and *Cldn3* showed strong temporal correlations with *Per1* and *Rev-Erbβ* expression, respectively (**Fig. S9C**). However, their circadian power varied across epochs: *Cldn1* was circadian from ∼E12.5, *Cldn3* from E10.5 to E14.5, and others (*Cldn2/11*, *Ocln*) did not exhibit strong rhythms until E15.5 (**Fig. 4F**, **Fig. S9D**). Similar to the core clock genes, the developmental timeline of circadian power in tight junction transcripts clustered independently of their promoter motifs (e.g., *Cldn1/3* and *Ocln* did not cluster together despite the presence of validated E-boxes) (**Fig. 4E, Fig. S9B**). If these transcripts translate to rhythmic protein levels, it would imply that the 4VCP can functionally gate the blood-CSF barrier circadianly as early as ∼E12.5, with CLDN3 acting as the first circadian gatekeeper, later relayed by CLDN1. Tight junction-mediated paracellular transport is a key pathway through which the blood-CSF barrier regulates the CSF (Ghersi-Egea et al., 2018).

Gene expression phases from E9 to E16 revealed a progressive maturation of the molecular circuit (**Fig. 4G**): phases were broadly distributed at E10.5, shifted dramatically into the complementary 12-h window by E14.5, and underwent strict bipartite polarization by E16.5, with *Clock*, *Chrono*, and *Cry2* antiphasic to the main group. This organization approaches the canonical TTFL phase organization but remains immature, as *Bmal1* is still in-phase with the E-box-driven group.

These patterns confirm three epochs of clock development: the undifferentiated epoch (E9.5-E12), with weak, non-canonical oscillations; the transition epoch (E12-E15), with period instability and phase reorganization; and the differentiated epoch (E15 onward), with stable, highly circadian oscillators approaching the phase organization of the mature TTFL.

### Maternal clock instability and feto-maternal reorganization

The absence of sustained *Bmal1-ELuc* rhythms in the embryonic 4VCP between E10.5 and E12.5 *ex vivo* (**Fig. 2B**), despite clear day-night variation *in vivo* over the same interval (**Fig. S8**), suggests that early circadian gene expression depends on maternal cues (Wong et al., 2022). Yet direct systemic maternal influence via the bloodstream is absent at this early stage. We performed spectral profiling of clock gene expression in the 4VCP of 102 wild-type dams (**Fig. S10A**). While all genes exhibited circadian rhythmicity at the transcript level (**Fig. 5A,B**; **Fig. S10A**), the circadian power, period, and phase of *Per2*, *Rora*, and *Rorc* decreased or fluctuated significantly as pregnancy progressed (**Fig. 5B,C**).

Comparison with non-pregnant controls revealed that pregnancy drastically altered both the amplitude and phase of clock transcripts, with the most profound shift at G12.5-G13.5, producing a distinct phase organization in the maternal 4VCP (**Fig. 5D**, **Fig. S10B,C**). Behavioral monitoring of free-running locomotor activity confirmed a significant drop in circadian power during pregnancy (\**p* < 0.05; two-sample *t*-test; **Fig. 5E,F**) consistent with reported declines in wheel-running during gestation (Yaw et al., 2021). In cases of presumed resorption, robust circadian rhythmicity was restored rapidly, accompanied by the return of the estrous cycle rhythm (**Fig. 5E**). These data demonstrate that the maternal circadian rhythm is transiently disrupted during pregnancy at both molecular and behavioral levels.

Pairwise correlations between dam and embryo clock gene expression revealed a global loss of maternal-fetal correlation at E12.5, contrasting with highly organized patterns at E16.5 (**Fig. 5G**, **Fig. S11A,B**). Maternal-embryonic coupling undergoes a transient destabilization around E12-E13 before re-establishing tighter correlation patterns after E14. Dual-radius phase plots revealed that correlation shifts were rarely one-sided (with *Bmal1* as a notable exception) (**Fig. 5H**), and the shift in maternal *Per2* phase coincided precisely with the reorganization of coupling patterns (**Fig. S11C,D**).

Finally, since both 4VCP clocks are destabilized during mid-pregnancy, we examined whether their physical interface, the placenta, was similarly affected. As the placenta matures, embryonic nutrient supply shifts from histiotrophic (yolk sac-based) to hemotrophic (blood-based), beginning around E10 and becoming fully functional by E12.5-E13.5 (Elmore et al., 2022). We separated embryonic and maternal placental tissues and recorded PER2::LUC and *Bmal1-ELuc* bioluminescence *ex vivo* from E9.5 to E20.5. The embryonic placenta lacked autonomous circadian oscillations (Čečmanová et al., 2019) (**Fig. S12A**), whereas the maternal placenta exhibited robust rhythms that underwent transient attenuation during mid-gestation (E13-E15) (**Fig. S12B**). Inverted Gaussian fits estimated a mid-gestational nadir in circadian power for PER2::LUC and *Bmal1-ELuc* at ∼20-30%, centered at E15.3 ± 0.4 and E15.6 ± 0.6, respectively. This circadian dip aligns closely with the functional maturation of the hemotrophic interface (Elmore et al., 2022).

Taken together, these results demonstrate that coupling between maternal and embryonic clocks is highly dynamic, undergoing shared transient disorganization around E13.5 before the embryonic system consolidates its circadian architecture.

## Discussion

Our study identifies the embryonic CP as the earliest detectable brain circadian clock, emerging ∼2.5 days before the SCN. We observed a staged assembly of the transcriptional clock circuit that coincides with cellular differentiation. The emergent rhythm exhibits hallmarks of a SNIC bifurcation, rendering the nascent clock highly sensitive to low-amplitude stimuli such as maternal core body temperature. Circadian profiling across embryonic development revealed three epochs characterizing *in vivo* clock emergence that closely track the differentiation program of the 4VCP. Strikingly, the middle transition epoch exhibits a transient destabilization of both maternal and embryonic 4VCP clocks (**Fig. S13**).

Contrary to the prevailing assumption that the SCN is the first circadian structure in the mammalian embryo, our data show that the 4VCP clock emerges significantly earlier. While our survey was limited to selected sagittal sections rather than the full three-dimensional extent of the brain, the 4VCP was consistently the earliest structure to display autonomous circadian rhythmicity. Of note, the first three structures identified as circadian by E15.5, the CP, SCN, and pituitary gland, are positioned to regulate brain homeostasis and endocrine signaling, and they maintain strong oscillations postnatally (Myung et al., 2018).

As the first clock in the embryonic brain, the 4VCP carries profound functional implications. Because the adult CP is a robust oscillator capable of entraining the SCN both *ex vivo* and *in vivo* (Myung et al., 2018), the early 4VCP clock could similarly influence the subsequent emergence of the SCN clock. Furthermore, as a secretory tissue, a circadian CP could rhythmically regulate CSF volume and composition, periodically releasing morphogens and growth factors (Kaiser and Bryja, 2020; Saunders et al., 2023; Lun et al., 2015). The neuroepithelium lining the ventricles would thereby receive temporally patterned cues capable of modulating transcription factor expression in neural progenitors and influencing cell fate decisions (Lamus et al., 2020; Lehtinen et al., 2011; Subashini et al., 2017). For example, oscillations of basic-helix-loop-helix (bHLH) factors (e.g., *Hes1*, *Ascl1*, *Olig2*) promote cell proliferation, whereas sustained expression drives differentiation (Imayoshi et al., 2013). By temporally gating differentiation, the progenitor pool can expand sufficiently before neurogenesis begins, a prerequisite for establishing the correct number of neurons and neural progenitors in the adult brain (Caviness et al., 1995; Kriegstein and Alvarez-Buylla, 2009).

Studying the embryonic 4VCP clock both *in vivo* (with maternal entrainment) and *ex vivo* (isolated) allowed us to map the relationship between clock assembly and the tissue differentiation program through three epochs. During the initial undifferentiated stage (E9.5-E11.5), despite a progenitor profile dominated by markers such as *Gli3* and *Ki67* (**Fig. 2E**), we observed early circadian expression of select clock genes (*Per2/3, Rora, Clock, Chrono*) *in vivo* (**Fig. S8**). This contrasts with *in vitro* studies showing that induced pluripotent stem cells (iPSCs), embryonic stem cells (ESCs), and other pluripotent stem cells (PSCs) lack circadian clocks (Yagita et al., 2010; Kowalska et al., 2010; Kaneko et al., 2023). This discrepancy likely arises because the *in vivo* embryonic 4VCP is exposed to maternally driven signals. Since the embryo lacks a direct blood connection to the dam at this stage, these signals must be ambient and low in amplitude. We demonstrated that the periodicity of maternal body temperature provides one such environmental cycle within the uterus, capable of driving the 4VCP clock until E16.5. Because this early ambient driving is restricted to highly responsive tissues such as the 4VCP, it underscores the importance of evaluating clock rhythmicity in a tissue-specific manner. Whole-embryo analyses, which dilute these localized rhythmic signals with non-rhythmic bulk, have previously missed these early clocks (Cao et al., 2022; Dolatshad et al., 2010).

During the transition phase (E12-E15), cellular differentiation is a prerequisite for full clock autonomy (Yagita, 2024; Yagita et al., 2010; Kowalska et al., 2010; Kaneko et al., 2023; Umemura et al., 2017). However, we note that *ex vivo*, the rhythm of the canonical activator *Bmal1* appears much later than that of *Per2*, suggesting that this embryonic clock is not initially established by classic BMAL1:CLOCK-driven feedback. *In vivo*, as differentiation begins, the embryonic rhythm is notably weakened, although transcripts like *Clock*, *Cry2*, and *Cldn3* remain circadian. This unstable epoch is mirrored in the maternal 4VCP and placenta, where we observed a significant decrease in PER2::LUC and *Bmal1-ELuc* circadian power, but not in maternal peripheral organs like the liver (Canaple et al., 2018). The mid-gestational instability coincides with increased activity of 11β-hydroxysteroid dehydrogenase type 2 (11β-HSD2; E12.5-E16.5), which inactivates glucocorticoids (Salvante et al., 2017). As the placenta matures, this selective endocrine gating likely attenuates maternal influence (Astiz and Oster, 2021), creating a shielded developmental window for the embryonic clock to undergo autonomous self-organization.

The discrepancy between our *in vivo* and *ex vivo Per2* data during this epoch warrants careful interpretation. One possibility, proposed by others (Yagita, 2024), is that the loss of circadian rhythmicity in many transcripts around E12-E15 reflects active suppression of the embryonic clock to permit differentiation. Since the 4VCP clock is autonomous *ex vivo* by E12.5, fluctuating maternal signals, which are themselves unstable during this window, could exert this inhibitory effect on the embryonic clock. An alternative interpretation is that the period instability during the transition epoch (E12-E14.5) reflects a transient state of low intra-tissue synchrony rather than active suppression. Before E12, nascent 4VCP oscillators near the SNIC threshold possess intrinsically variable periods but are frequency-locked by maternal temperature cycles. Around E12-E13, the maternal clock destabilizes. Compounded by progressive placental shielding, the maternal drive withdraws before endogenous intercellular coupling has matured, even as the underlying cellular bifurcation dynamics shifts toward a Hopf regime during TTFL maturation. Consequently, the tissue-level readout averages over desynchronized cells, effectively blunting any underlying rhythms. By E14.5, intrinsic coupling rises abruptly, and individual cellular rhythms become coherently detectable in whole-tissue *in vivo* quantification. Beyond the effects of entrainment or synchrony, the apparent delay in *Per2* rhythmicity may simply reflect the different biomolecules being measured. At E11.7, PER2 protein oscillates with a ∼24 h period *ex vivo*, whereas *Per2* transcripts do not yet cycle *in vivo*. This discrepancy is expected, as robustly cycling proteins frequently originate from non-cycling transcripts through post-transcriptional regulation (Qian et al., 2023; Robles et al., 2014; Mauvoisin et al., 2014). Conversely, BMAL1 protein can be post-translationally modified to remain functionally inactive, potentially preventing premature interference with ongoing developmental programs (Umemura et al., 2022). This illustrates why transcript presence does not always equate to functional integration into the TTFL circuit.

Following this transient destabilization, the fully differentiated 4VCP (E15-E16.5) exhibits a more robust, autonomous clock *ex vivo*. *In vivo*, the majority of embryonic clock genes regain circadian rhythmicity and organize their phases into two antiphasic groups, with *Clock* adopting the opposing phase. This restoration likely reflects the combined influence of SCN emergence, strengthened intercellular synchrony, and the intrinsic capacity of *Bmal1* to oscillate. In the dam, although the 4VCP clock appears restored, the free-running behavioral rhythm remains impaired, potentially reflecting continued instability in the SCN. By contrast, in the 4VCP, both maternal and embryonic genes align in phase during this epoch, suggesting a synchronized maternal-fetal relationship. However, our study design does not allow us to determine the directionality of maternal-embryonic coupling.

Multiple lines of evidence indicate a relatively autonomous embryonic clock by late gestation. In mice, maternal influence on pup behavioral synchrony occurs early rather than late in gestation (Davis and Gorski, 1988; Jud and Albrecht, 2006), and in humans, preterm infants display detectable circadian-like rhythms in body temperature, heart rate, and rest-activity in the NICU (Bueno and Menna-Barreto, 2016; Govindan et al., 2024), with 24 h sleep-wake rhythms appearing earlier in preterm than in full-term infants (Guyer et al., 2015). Our data show that the 4VCP remains sensitive to 0.5 °C temperature cycles at E16 but not E18, consistent with entrainment observed in late embryos following maternal restricted feeding (Canaple et al., 2018).

The early emergence of the circadian clock in the 4VCP at E11.5-E12.5 challenges the long-held view of the SCN as the origin of circadian pacemaking in the mammalian brain and, by extension, the presumed locus of developmental onset. Our findings further reveal that the clock emergence in the CP is a discrete, non-canonical developmental process, with rhythmic expression of *Rora*, *Per2/3*, and *Chrono* detectable as early as E9.5. Furthermore, while mature circadian oscillators are classically modelled by Hopf bifurcations, this early developmental transition is governed by highly sensitive SNIC bifurcation dynamics. The multi-stage assembly and sudden external emergence of the clock strongly support the paradigm that circadian clock onset is a programmed developmental event. By characterizing the precise timing, locus, and mechanism of the brain’s first clock, this work provides a new framework for understanding maternal-fetal interactions and insights into early circadian homeostasis with direct relevance to preterm and neonatal care.

## Material and Methods

### Mice

Homozygous PER2::LUC mice (B6.129S6-*Per2^tm1Jt^*/J, RRID: IMSR_JAX:006852) and heterozygous *Bmal1-ELuc* mice (Nakajima et al., 2010; a gift from Daisuke Ono, Nagoya University), maintained on the C57BL/6J background, were crossed with wild-type C57BL/6J animals to generate heterozygous PER2::LUC^+/-^ or *Bmal1-ELuc*^+/-^ embryos. Mice were maintained under equinox light-dark (LD) cycles (12h light:12h dark; lights on at 07:00, off at 19:00), with schedules controlled by LocoBox software (Truong and Myung, 2023). Ambient temperature and humidity were maintained at 24.0 ± 1.8 °C; 50.2 ± 8.7% RH. Zeitgeber Time (ZT) 0 was defined as the time of lights-on. For tissue sampling at ZT14 and ZT20, a reversed dark-light (DL) cycle was used (lights on at 22:00, off at 10:00) (**Fig. S1A**). We validated that embryonic gene expression levels under the reversed cycle matched those under standard LD cycles at E13.5 (**Fig. S1B**). All animal protocols were approved by the Institutional Animal Care and Use Committee (IACUC) of the Taipei Medical University Laboratory Animal Center (protocols LAC-2019-0430, LAC2022-0381, LAC2022-0477, LAC2023-0037, and LAC2023-0038).

### Circadian actimetry

Female wild-type C57BL/6J mice (6 weeks old) were housed in constant darkness (DD) for two weeks to get their baseline activity, followed by one week under a 12:12 LD cycle to synchronize their phase with the LD-housed males. Following a single night of mating, females were isolated, weighed, and returned to DD for ∼21 days. Locomotor activity was continuously recorded using LocoBox (Truong and Myung, 2023). Pregnancy was confirmed by weighing females at G8.5 and/or G13.5 under DD, and activity monitoring continued until parturition. Cage changes were restricted to G8.5 or G13.5 to minimize disruptions to locomotor recordings.

### Tissue isolation and explant culture

#### Direct dissection

Tissue culture and bioluminescence recording were performed according to our previous protocols (Myung et al., 2018). Pregnant female mice were anesthetized with isoflurane and euthanized at defined ZTs. Brains and embryos were rapidly extracted into ice-cold Hank’s balanced salt solution (HBSS; Sigma-Aldrich) supplemented with penicillin-streptomycin (10,000 U/mL; Gibco) and HEPES (1 M, pH 7.4). Embryonic tissues were dissected under a stereomicroscope after decapitation. For E9.5, the prosencephalon (containing SCN and LVCP progenitors) and rhombencephalon (containing 4VCP progenitors) were collected. For E10.5-E12.5, the 4VCP was extracted from the inferior part of the hindbrain roof plate epithelium (lower rhombic lip, rhombomere 2-8), located between the otic vesicles and the posterior part of rhombomere 1 (Hunter and Dymecki, 2007). The cortical midline served as the anatomical landmark for LVCP progenitors. After E12.5, the CP dissection was similar to that of the adult. The precision of 4VCP isolation throughout development was validated by RT-qPCR (**Fig. S2**). SCN progenitors from E10.5 to E13.5 were dissected from the ventral hypothalamus between the optic cups. After E13.5, the SCN and the optic chiasm were clearly distinguishable, and dissection was performed as in the adult. This protocol was also used to isolate maternal and embryonic tissues in ice-cold PBS for biomolecular experiments.

#### Placenta dissection

The maternal decidua (light pink outer layer, representing maternal placenta) was mechanically separated from the junctional zone and labyrinth (darker pink inner layer, representing the embryonic placenta). The loose junction between these two layers allowed dissection with forceps (Čečmanová et al., 2019).

#### Tissue slicing and explant culture

Embryonic heads or brains (E11.5, E13.5, E15.5) were briefly submerged in two successive baths of low-melting-point agarose (2% w/v; CAS 9012-36-6) at 37 °C, embedded in an agarose block, and sliced in ice-cold HBSS using a vibratome (VT1000S, Leica Biosystems). Slices (300 µm thick) were cut using a diamond blade (Delaware Diamond Knives) advancing at ∼0.025 mm/s. Explants were cultured on culture membrane inserts (PICM0RG50, Merck Millipore) in 35-mm dishes containing 1 mL DMEM (Sigma-Aldrich) supplemented with B-27 (Gibco) and 200 µM beetle luciferin (Promega). Dishes were sealed with high-vacuum grease (Molykote, DuPont) and maintained at a strictly controlled 37.00 ± 0.01 °C in a hybrid refrigerator-heater incubator (MIR-154-PA, PHCbi), monitored closely by placing an iButton (DS1921L, Maxim Integrated) adjacent to the dishes.

#### iDISCO tissue clearing and immunofluorescence labeling

Post-culture brain slices were cleared using the standard iDISCO protocol (https://idisco.info/idisco-protocol). Slices were fixed in 4% PFA, washed in PBS for 3 days, dehydrated in a MeOH/H_2_O series, and incubated overnight in 66% DCM/33% MeOH. Tissues were bleached in 5% H_2_O_2_ overnight and rehydrated. Immunolabeling was performed at 37 °C, beginning with permeabilization (1 day) and blocking (1 day), followed by primary antibody incubation against TTR (chicken, 1:500, Genetex GTX85112; AB_10723946) for 2 days. Slices were washed and incubated with a secondary antibody (anti-chicken Alexa Fluor 568, 1:500, Abcam ab175477; AB_3076392) for 2 days. Following subsequent washing and blocking, slices were incubated with a conjugated RORA antibody (1:500, Bioss BS-17164R-A350). Slices were then transferred back to the micro glass vials to undergo final clearing, beginning with dehydration in MeOH/H_2_O followed by a 3h incubation in 66% DCM/33% MeOH at room temperature. Finally, two 15-min incubations in 100% DCM were performed before mounting the slices in dibenzyl ether under sealed coverslips, and imaged using a DM IL fluorescence microscope (Leica Microsystems).

### Time-lapse bioluminescence luminometry and imaging

#### Whole-tissue luminometry

PER2::LUC fusion protein reporters tracked PER2 abundance through bioluminescence (RRID: IMSR_JAX:006852), while *Bmal1-ELuc* reporters tracked *Bmal1* transcriptional activity (Nakajima et al., 2010). The overall bioluminescence from each tissue was recorded every 15 min (108 s integration time per dish) using LumiCycle 32 (Actimetrics) with four photomultiplier tubes (PMTs), housed in a refrigerator-heater incubator for precise temperature control (MIR-154-PA, PHCbi). Temperature was recorded continuously throughout the experiment.

#### Single-cell imaging

Imaging was conducted on a custom incubator-microscope system equipped with a cooled-CCD camera (Orca R2, Hamamatsu) with water cooling, combined with a 4x objective (NA 0.16, UPLXAPO, Olympus) coupled to a 0.35x reduction lens (U-TV0.35XC-2, Olympus), and a stage-top incubator (INUG2-WELS, Tokai Hit). Images were acquired hourly (60 min exposure time) with 4×4 binning using Micro-Manager 1.4 (Vale Lab, UCSF) via the DCAM-API (Hamamatsu) driver.

### Real-time quantitative PCR (RT-qPCR)

#### RNA Isolation, cDNA synthesis, and RT-qPCR

Embryonic or maternal 4VCPs collected at ZT2, ZT8, ZT14, and ZT20 were pooled by litter (3-11 embryos/litter, minimum 2-3 litters per time point). Total RNA isolation and RT-qPCR followed our previous protocol (Myung et al., 2018). 300 µL of TRK lysis buffer containing 2% β-mercaptoethanol was added to each sample, followed by an equal volume of chloroform after intensive homogenization. After phase separation and ethanol addition, microcolumns from the ENZA MicroElute Total RNA Kit (Omega Bio-Tek) were used to collect total RNA, with a final elution volume of 21 µL in DEPC-treated water. After evaluating the RNA concentration, cDNA was synthesized from 0.5-1 µg of total RNA using SuperScript IV Reverse Transcriptase (Thermo Fisher Scientific) with oligo(dT)_20_ primers. PowerUp SYBR Green Master Mix (Applied Biosystems) was added to the cDNA samples, and RT-qPCR was performed in technical triplicate on a QuantStudio 3 Real-Time PCR System (Thermo Fisher Scientific). Primer sequences are detailed in **Table S1**.

#### Normalization

*Gapdh*, *Actb* (*β-actin*), and *Rpl13a* served as internal controls for embryonic samples (Hildyard et al., 2022). G*apdh* and *Actb* were used for maternal samples. After outlier exclusion (ROUT test, Q = 1%), the relative mRNA abundance for a given embryonic stage (*n* > 8) was calculated by normalizing to the maximum abundance observed between E9.5 and E16.5. For example, at E9.5:

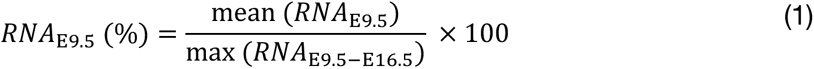

### Spectral analysis of bioluminescent time series and time of emergence quantification

#### Preprocessing

Visualization and preprocessing of bioluminescence oscillations, including periodogram and fast Fourier transform (FFT)-based spectral analysis, were performed on Mathematica 13 (Wolfram Research) using the custom PMTAnalysis package (http://sourceforge.net/projects/imaginganalysis). The first 6 h of all recordings were truncated to remove handling transients. Circadian periods were estimated using FFT, and amplitudes were quantified as the root mean square (RMS) of detrended oscillations over fixed windows (7 days for luminometry/4VCP imaging; 4 days for whole-brain slice imaging). For image analysis of brain slices or the 4VCP, a 2×2-pixel grid was applied to the timelapse images after removal of outlier pixels, and each ROI’s standard deviation, center of mass, and mean grey value were measured using ImageJ (NIH).

#### Circadian power

Using a sliding-window FFT (2-day for luminometry, 3-day for imaging), evolution of spectral power was tracked and visualized as spectrograms over time. Circadian power for each time window was calculated by integrating the spectral power within the circadian band (24 ± 4 h) relative to the full period range (3-58 h) (**Fig. S3**). This generated a time series reflecting the robustness of the circadian rhythm that accounts for damping, period variability, and noise. In addition, the overall circadian power was computed as the average of these window-wise circadian power values across the entire recording. For a perfect sinusoid in a 2-day window, the maximal circadian power reached ∼60%, whereas for transient, rapidly damped oscillations, it fell below 20% (**Fig. S3**). The blank control (luciferase medium without tissue explant) showed a maximal circadian power of ∼20% (*n* = 10) (**Fig. 2D**). For SCN and 4VCP explants, the half-maximal level of the fitted circadian power curve across development was ∼30% (30.83 ± 2.36% and 31.28 ± 1.70%, respectively).

#### Emergence Time (𝑻_𝒆_)

Nonlinear curve fitting using a sigmoid function was applied to rigorously identify the onset of PER2::LUC and *Bmal1-ELuc* circadian rhythms. Blank recordings were used to estimate baseline circadian power and to provide null data points prior to E9.5. 𝑻_𝒆_ was defined as the time point at which one-half of the maximum circadian power was reached:

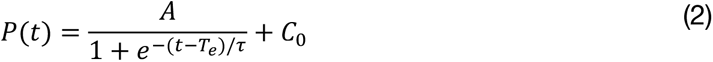

where 𝑃(𝑡) is the circadian power at time *t* (time of sampling for explant), 𝐴 is the amplitude above baseline, 𝜏 is the time constant, and 𝐶_0_ is the baseline determined from blank recordings.

### Circadian promoter motif analysis

We analyzed promoter-proximal regions (-3 kb to +2 kb relative to TSS in GRCm39) of 12 core circadian genes from our qPCR panel (**Table S1**). Gene coordinates and strand orientation were retrieved from Ensembl via the REST API (https://rest.ensembl.org/). To ensure all windows were extracted in the transcriptional orientation, minus-strand loci were reverse-complemented, so that upstream and downstream were always defined relative to the direction of transcription. We scanned these sequences for known transcription factor binding motifs relevant to circadian regulation, including canonical and variant E-boxes (CACGTG, CACGTT, CACGTA), D-boxes (TTATGAA, TTATGTA), REV-ERB/ROR response elements (RRE/RORE) (AGGTCA, GGGTCA, [A/U]GGTCA), glucocorticoid response elements (GREs) including canonical palindromic (AGAACA[ACGT]TGTTCT), *Per2*-specific variants ([AG]G[ACGT]TGT[TC]CT), as well as half-sites ([AG]G[TA]ACA, TGT[TC]CT), and cAMP response element (CRE) half-sites (CGTCA, TGACG). Motifs were represented either as exact sequences or regular expressions to capture degenerate variants. Both forward and reverse-complement strands were scanned. Motif hits were identified using regular expression search with overlapping match lookahead, followed by a priority-based deduplication scheme favoring full palindromes and canonical motifs over partial or variant sites. Motif density was expressed as the number of motifs per kilobase of sequence for each gene and region. These densities are visualized as normalized motif densities (**Fig. S7B, S9B**).

### Hierarchical clustering and spectral analysis of gene expression profiles

#### Hierarchical clustering

Expression levels were normalized using the internal control genes and standardized using Z-scores. Pairwise distances were calculated using the Pearson correlation 𝑟*_ij_* as the distance metric between expression of genes 𝑖 and 𝑗, and hierarchical clustering was implemented via the seaborn.clustermap function (v0.11.2) in Python (v3.6.13). Clustering was performed using average linkage, and results were visualized as a cluster heatmap with a dendrogram that represents the similarity relationships. Average linkage distance 𝐷(𝐴, 𝐵) was calculated as:

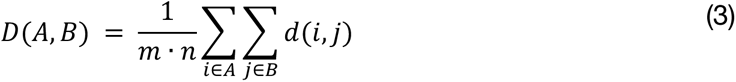

where 𝑚 and 𝑛 are the numbers of genes in clusters 𝐴 and 𝐵, and 𝑑(𝑖, 𝑗) is the distance (1-𝑟*_ij_*) between genes 𝑖 and 𝑗. We identified representative genes for each cluster by cluster centroids calculated by averaging the Z-scored profiles of all member genes across time points. The gene with the highest Pearson correlation to the centroid was designated as the representative, as it best reflected the average temporal pattern of its cluster.

#### Synchrosqueezed continuous wavelet transform (SST-CWT)

To analyze sparsely sampled, non-stationary embryonic gene profiles, we applied SST-CWT to sharpen time-frequency estimates, which reassigns CWT coefficients from a range of supporting frequencies to a more precise instantaneous frequency (Daubechies et al., 2011). We implemented it using the Python cwt_sst function from the ssqueezepy library (v0.6.5; https://github.com/OverLordGoldDragon/ssqueezepy). Normalized gene expression time series were detrended, mirrored, and extended prior to transform using a Morlet wavelet (90 voices per octave). Although SST-CWT yields an amplitude spectrum, we refer to it here as a “power spectrum” (the true power is the squared amplitude). Circadian power was calculated identically to the bioluminescence data (24 ± 4 h band relative to the 3-58 h full range).

#### Phase dynamics and visualization

We estimated instantaneous phase (radians) along ridges of the time-frequency representation. Instead of plotting phase alone, we generated phase-weighted spectrograms, which multiply SST-CWT amplitude by its corresponding phase angle to highlight dominant oscillatory components with phase context. To improve visual clarity, colormap normalization and adaptive contrast scaling were applied using the maximum absolute value of each map. An illustration of this approach using simulated data is shown in **Fig. S7A**.

#### Circular phase plots

To track phase across development (E10.5, E12.5, E14.5, E16.5), we constructed circular phase plots based on the estimated instantaneous phase within the circadian range (20-28 h). For each embryonic cluster, the representative gene was plotted clockwise in a phase plot resembling a conventional clock, with 0 rad positioned at 12 o’clock. Nonrepresentative genes from the same cluster were plotted with smaller, semi-transparent markers for comparison. The left semicircle was shaded to distinguish early vs late phases. Marker shape and color indicated cluster identity, and genes with circadian power ≥ 20% were outlined in black. To assess potential maternal-embryonic coordination, dual-radius phase plots were used: the inner ring showed embryonic cluster representatives, and the outer ring showed selected maternal genes. Maternal-embryonic pairs were connected with dotted lines when both exhibited circadian power ≥ 20% and their phase difference was < *π*/2 rad (≤ 6 h), indicating possible phase alignment.

#### Sliding-window correlation

To assess temporal coordination among maternal and embryonic genes, time series were padded by duplicating the first and last two points, and 5-timepoint sliding windows were applied. Pearson correlation matrices were computed for each window, generating a time-resolved sequence of correlations. Examples of dynamic correlation are shown in **Fig. S11A**.

#### Normalized spectral entropy and identification of transition epoch

To quantify the periodicity complexity in each detrended bioluminescence trace, we calculated the spectral entropy from the SST-CWT of each gene’s time series as:

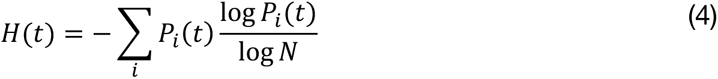

where 𝑃*_i_*(𝑡) is the normalized power in the 𝑖-th frequency bin at time 𝑡, and 𝑁 is the number of frequency bins. Low entropy corresponds to power concentrated in a single frequency, while high entropy indicates power dispersed across multiple frequencies. To define the transition epoch without relying on an arbitrary threshold, we employed a permutation clustering approach on the pooled embryonic clock gene entropy trajectories. Mean normalized spectral entropy across all genes was computed at each time point, and contiguous time points exceeding the median were identified as candidate clusters. To assess significance, we generated a null distribution of maximum cluster sizes by independently permuting each gene’s entropy trace 1,000 times. Clusters whose size exceeded the 95th percentile of the null distribution were considered significant transition epochs and visualized as shaded intervals on the mean entropy trajectory.

### Statistical Analysis

Data are reported as mean ± standard error of the mean (SEM) unless otherwise specified. Circular phases are presented as mean ± circular standard deviation (SD). For parametric curve fits, we report best fit estimate ± standard error (SE), obtained from the asymptotic covariance matrix of the nonlinear least-squares fit. Statistical comparisons used unpaired or one-sample *t*-tests for two-group comparisons, the Mann-Whitney U test for non-parametric comparisons (**Fig. S1**), one-way ANOVA with Tukey’s post hoc test for multiple groups, and two-way ANOVA followed by the appropriate post hoc test for interactions over time. Linear relationships were assessed using ordinary least-squares regression with the coefficient of determination (*R*^2^) as goodness of fit, and a two-sided *p* value. For nonlinear fits, model selection was guided by the Akaike Information Criterion (AIC) and the Bayesian Information Criterion (BIC). For temperature entrainment experiments (**Fig. 3F**), phase drift relative to zero was assessed using two one-sided tests for equivalence (TOST; equivalence bounds ±0.483 rad/cycle) together with one-sample *t*-tests. At E14, 4VCP drift was not statistically different from zero (-0.154 ± 0.109 rad/cycle, *p* = 0.254), whereas lung drift was statistically equivalent to zero (0.078 ± 0.025 rad/cycle, *p* = 0.053). At E16, both tissues showed drift equivalent to zero (4VCP: 0.174 ± 0.089 rad/cycle, *p* = 0.146; lung: 0.018 ± 0.010 rad/cycle, *p* = 0.137). At E18, the 4VCP displayed significant negative drift exceeding equivalence bounds (-0.473 ± 0.042 rad/cycle, *p* = 0.0015), whereas the lung remained equivalent to zero (0.068 ± 0.019 rad/cycle, *p* = 0.0396). No data were excluded as outliers except in the following cases: spurious FFT-derived period estimates in non-periodic time series, clear failures in RNA quantification, and dispersed amplitude or period values from 4VCP ROIs, where outliers were identified using the ROUT test (Q = 1%). All statistical analyses were performed in GraphPad Prism 9 (GraphPad) and Mathematica 13 (Wolfram Research).

## Acknowledgements

We thank Daisuke Ono (Nagoya University) for providing the *Bmal1-ELuc* mice and Lillian Byer for critical reading of the manuscript. This work was supported by the National Science and Technology Council (NSTC) (109-2314-B-038-071, 109-2320-B-038-020, 110-2314-B-038-162, 110-2311-B-038-003, 111-2314-B-038-008, 112-2314-B-038-063, 113-2314-B-038-121, 114-2320-B-038-052-MY3, 114-2811-B-038-015) and the Higher Education Sprout Project of the Ministry of Education (MOE) in Taiwan. Figures were created using BioRender.com.

## Data availability

The data supporting the findings of this study are available from the corresponding author upon reasonable request.

## Supplementary Materials

**Table S1.**
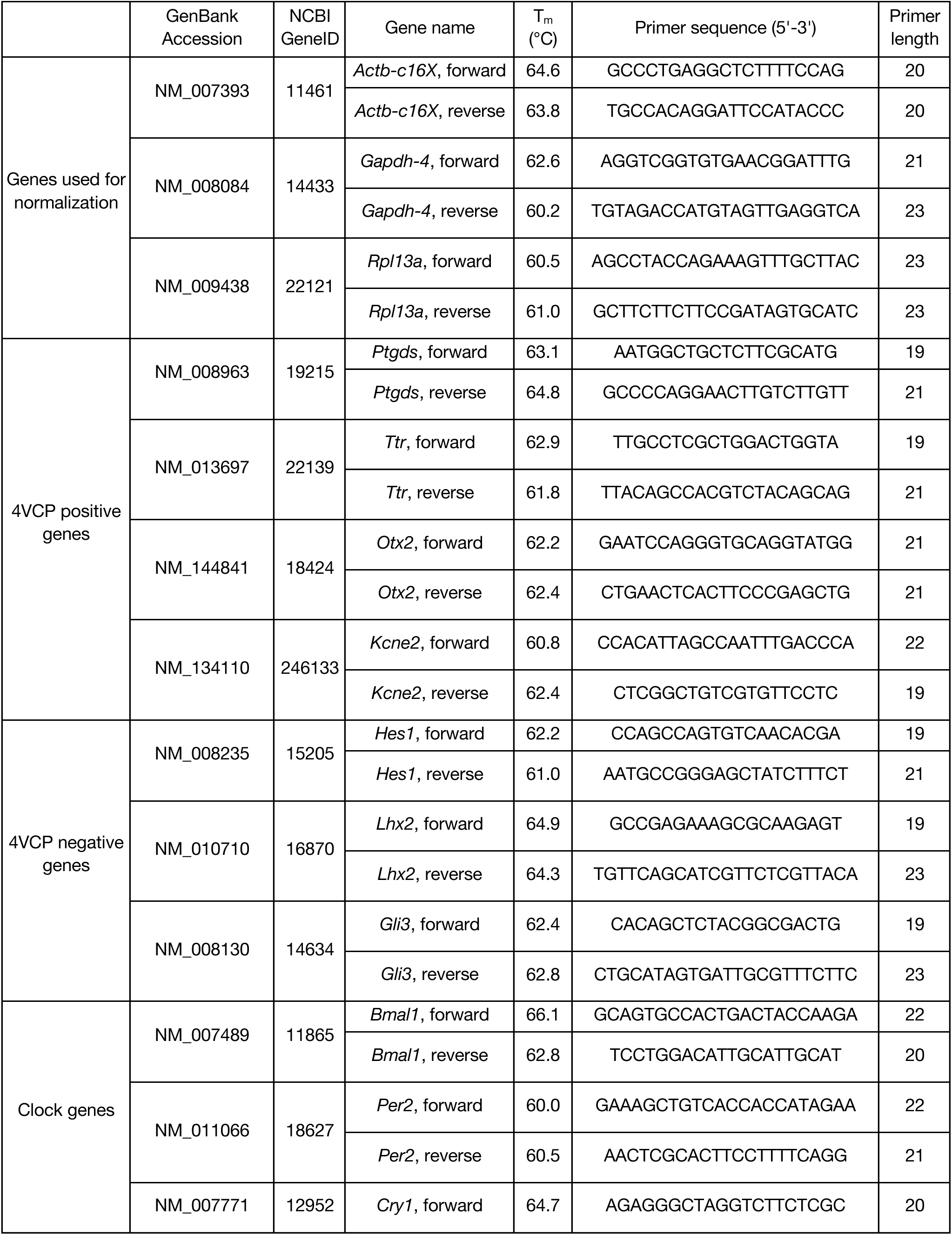
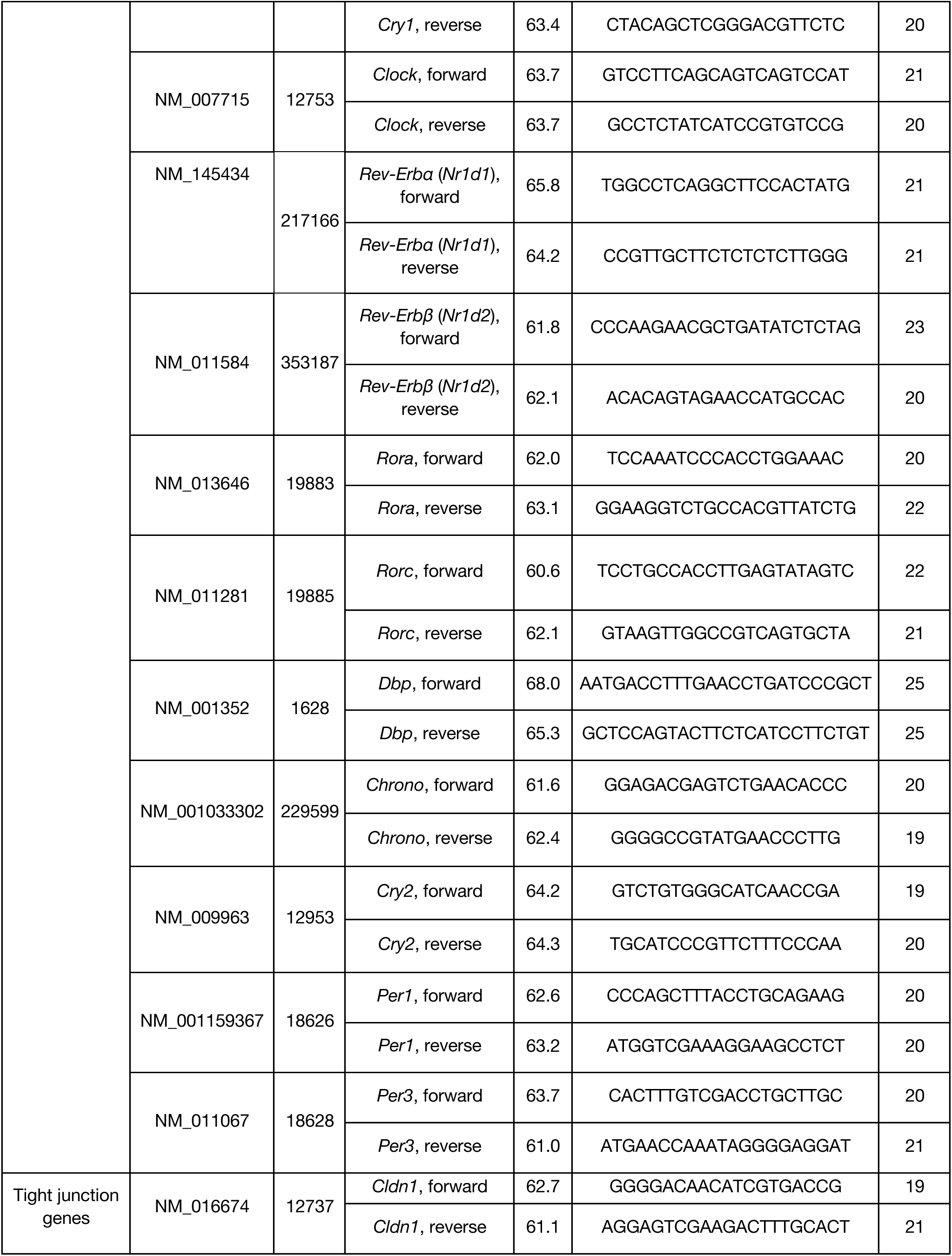
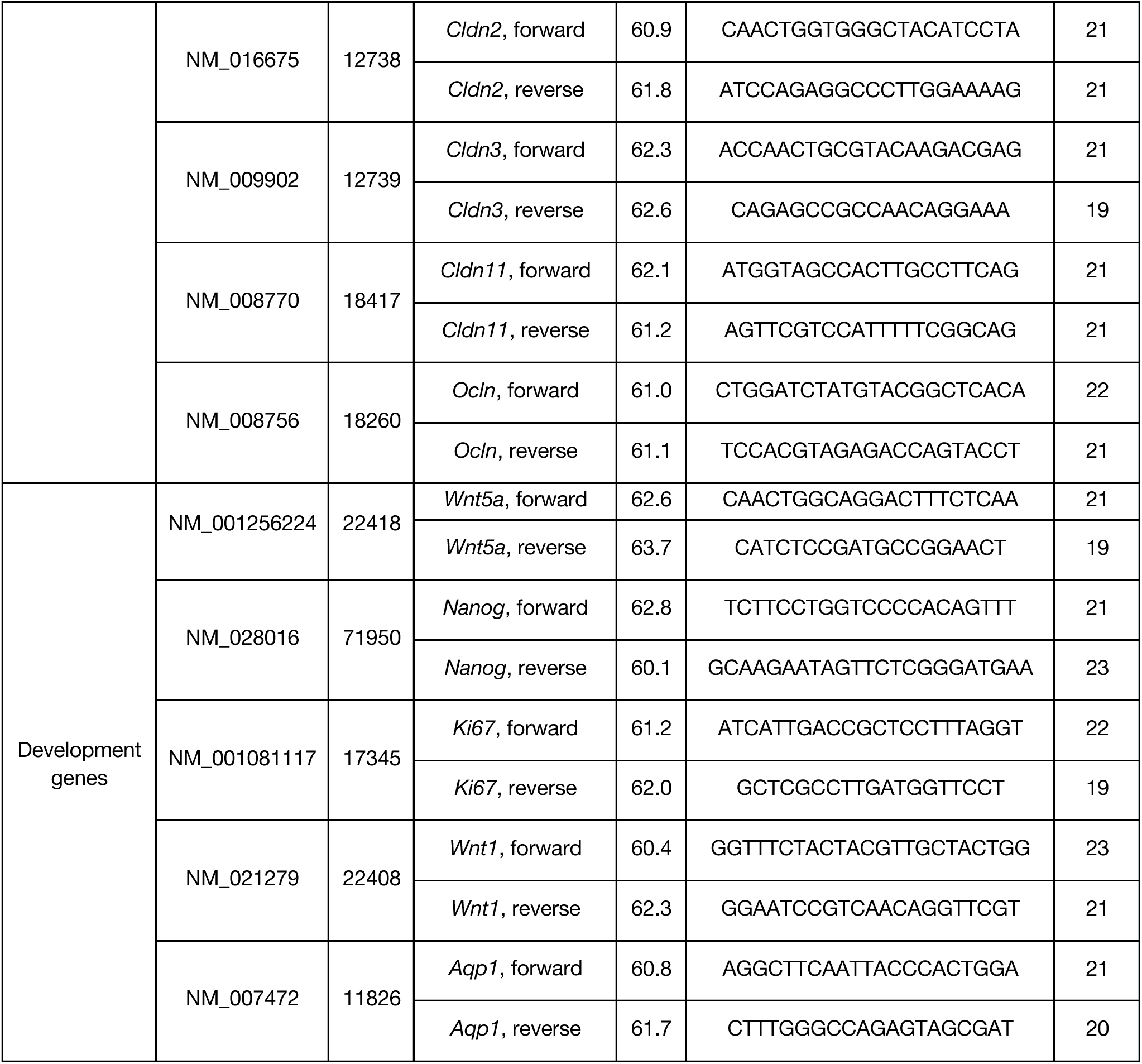
Primer sequences used for qPCR. *T_m_* indicates melting temperature.

**Table S2.**
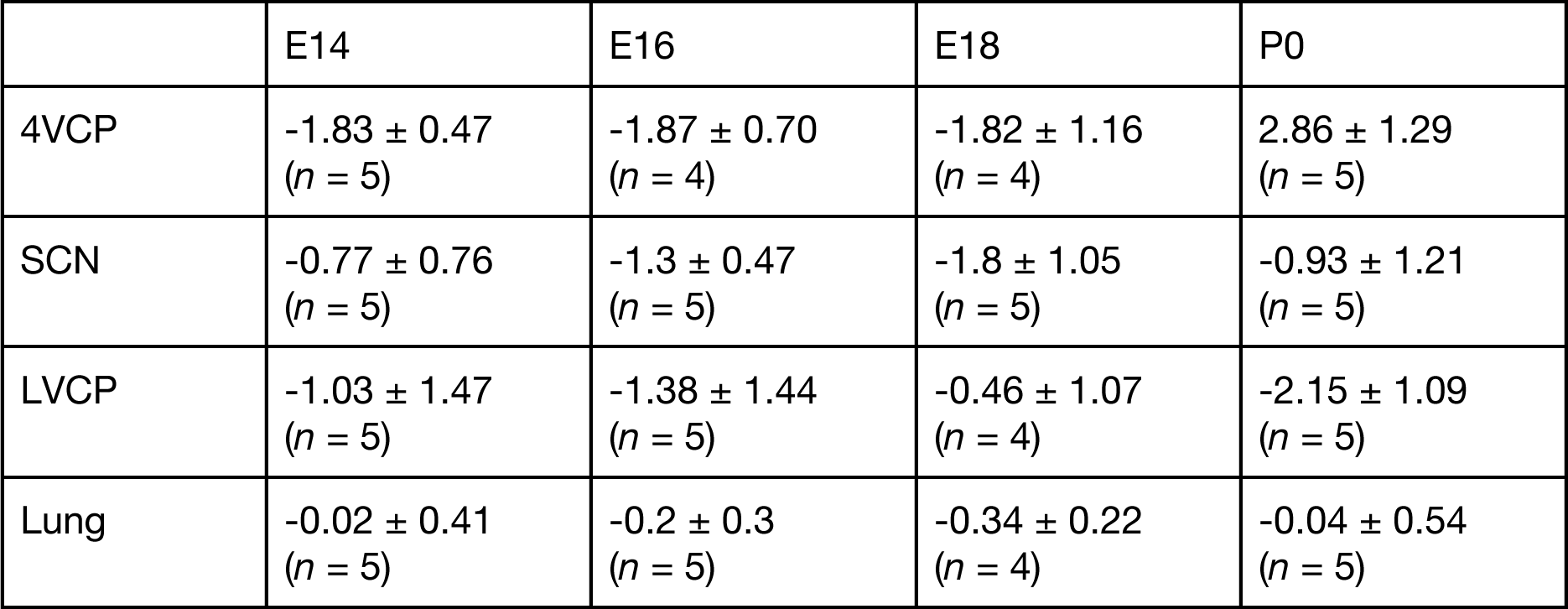
Phase of entrainment (rad) under a 26 h temperature cycle. Shown are circular mean ± circular SD from 4-5 explants. For each explant, peak phase was averaged over the final 7 days of recording under the 26 h cycle (0 rad defined as the onset of the 37.5 °C epoch; *n*: number of explants). Negative values indicate phase lead relative to 0 rad.

**Figure S1.**
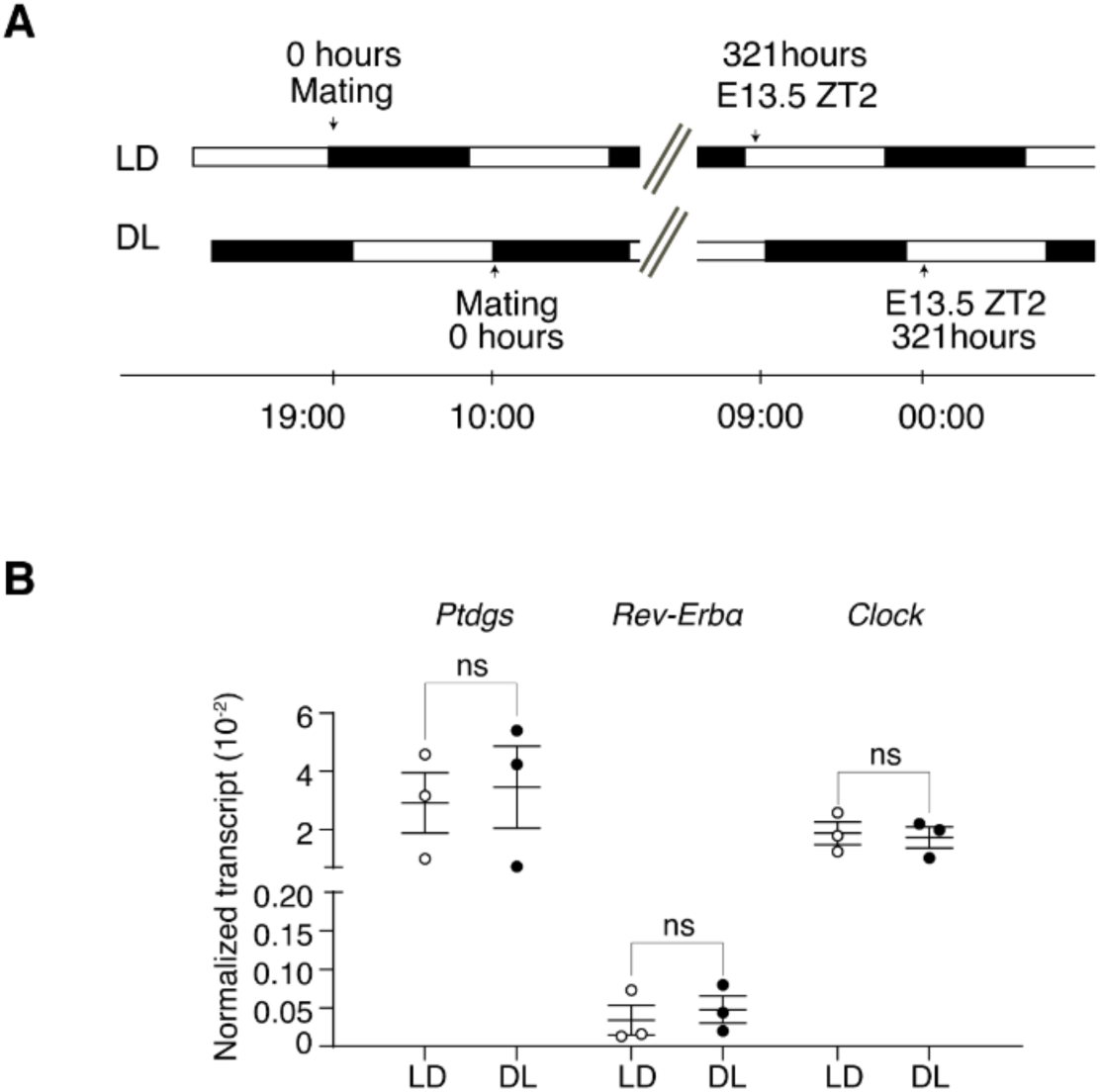
Comparison of mRNA expression between LD and DL protocols. (**A**) Scheme of the light-dark schedules for LD and DL (reversed-LD) conditions. (**B**) Normalized mRNA abundance of three representative genes (*Ptgds*, *Rev-erbα*, *Clock*) out of 22 tested, in the embryonic 4VCP sampled at the same ZT (ZT2) from 3 pregnant mice per condition. Mann-Whitney U test; ns, not significant. Error bars: mean ± SEM.

**Figure S2.**
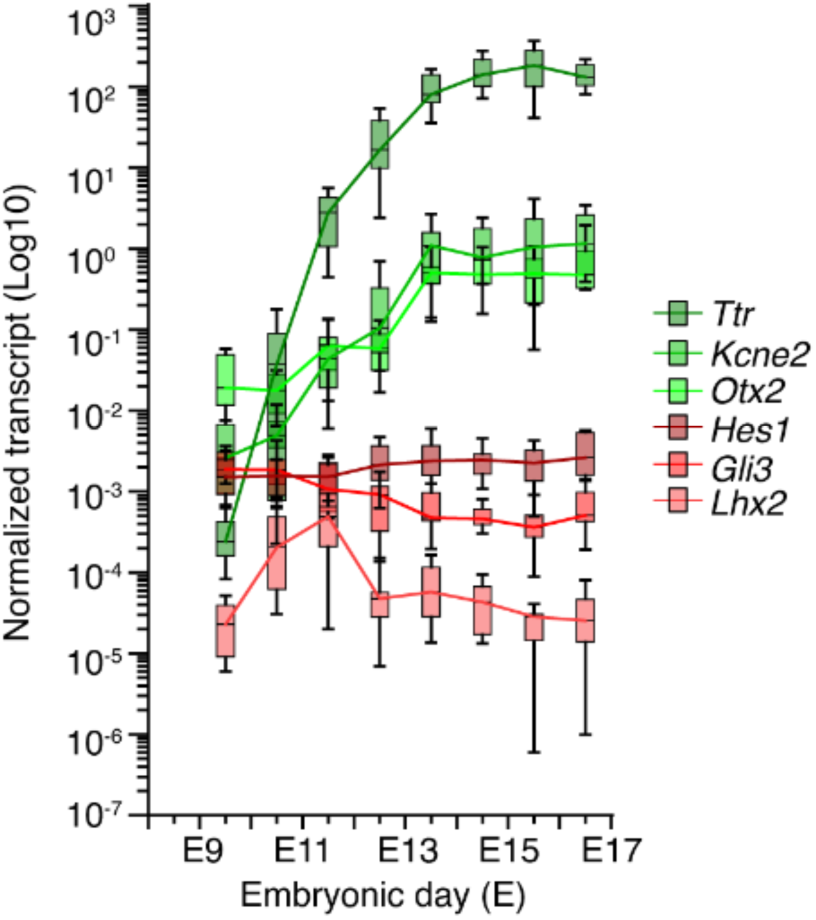
Molecular validation of 4VCP dissection across embryonic stages. Normalized mRNA abundance of selected 4VCP-positive (*Ttr*, *Kcne2*, *Otx2*; green) and 4VCP-negative (*Gli3*, *Lhx2*, *Hes1*; red) markers in the embryonic 4VCP (*n* ≥ 8 litters per stage). Data are plotted on a log_10_ scale; box-and-whisker plots show the 10th-90th percentiles.

**Figure S3.**
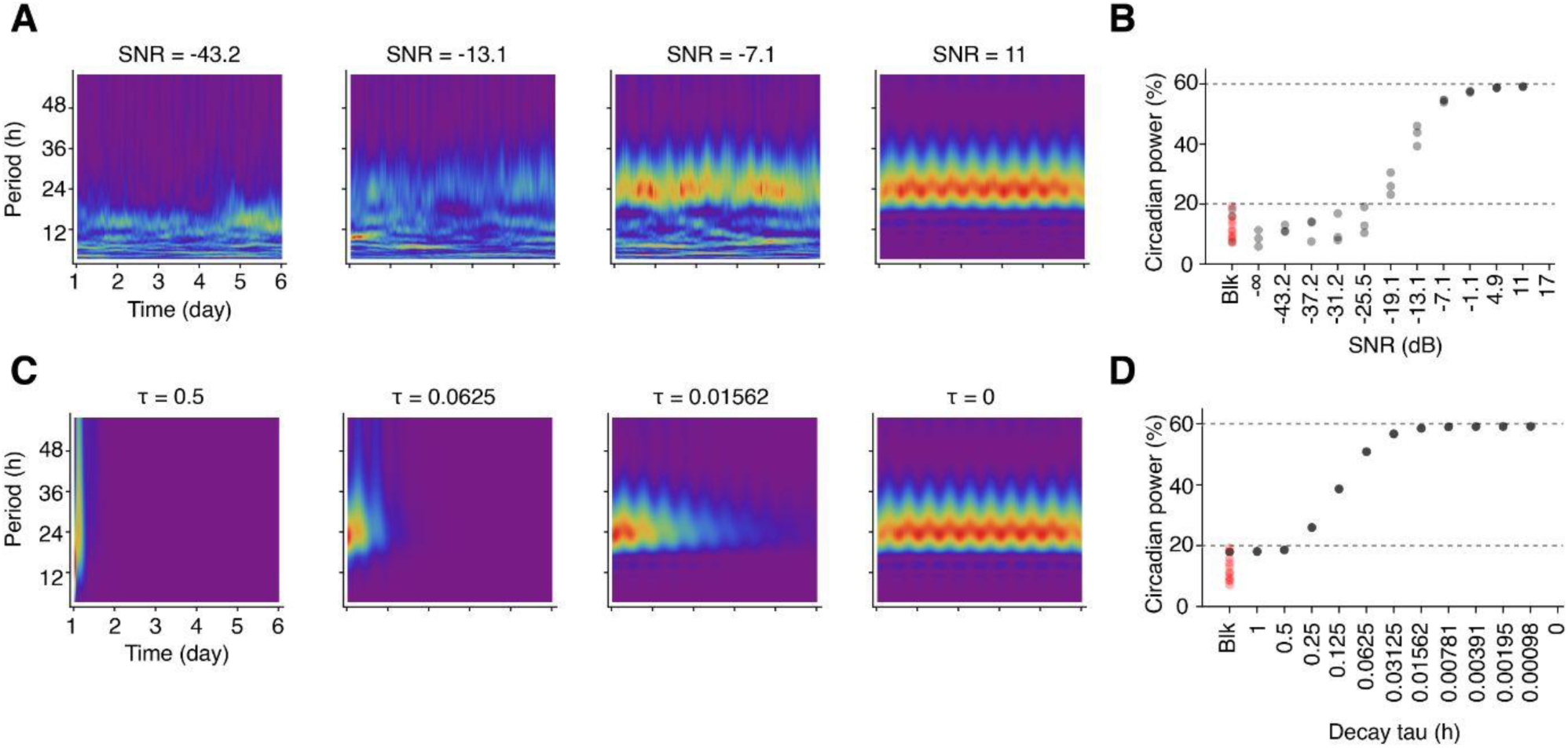
Quantification of circadian power in simulated bioluminescence time series. (**A**) Spectrograms from 2-day sliding-window FFT of simulated 24-h oscillations with varying noise levels, expressed as signal-to-noise ratio (SNR, dB). Increasing SNR progressively reveals the circadian peak at ∼24 h. (**B**) Circadian power as a function of SNR. Each point represents one simulation. Detectable circadian rhythms fall between 20% (detection threshold) and 60% (ceiling), indicated by dashed lines. Blank recordings (Blk) from actual experiments fall below 20%. (**C**) Spectrograms of simulated 24 h oscillations with varying damping time constants (*τ*). Shorter *τ* produces stronger damping and loss of rhythmicity. (**D**) Circadian power as a function of *τ*. Semi-transparent points represent individual simulations but overlap substantially. As in (B), circadian rhythms below the 20% threshold (dashed line) are considered non-detectable.

**Figure S4.**
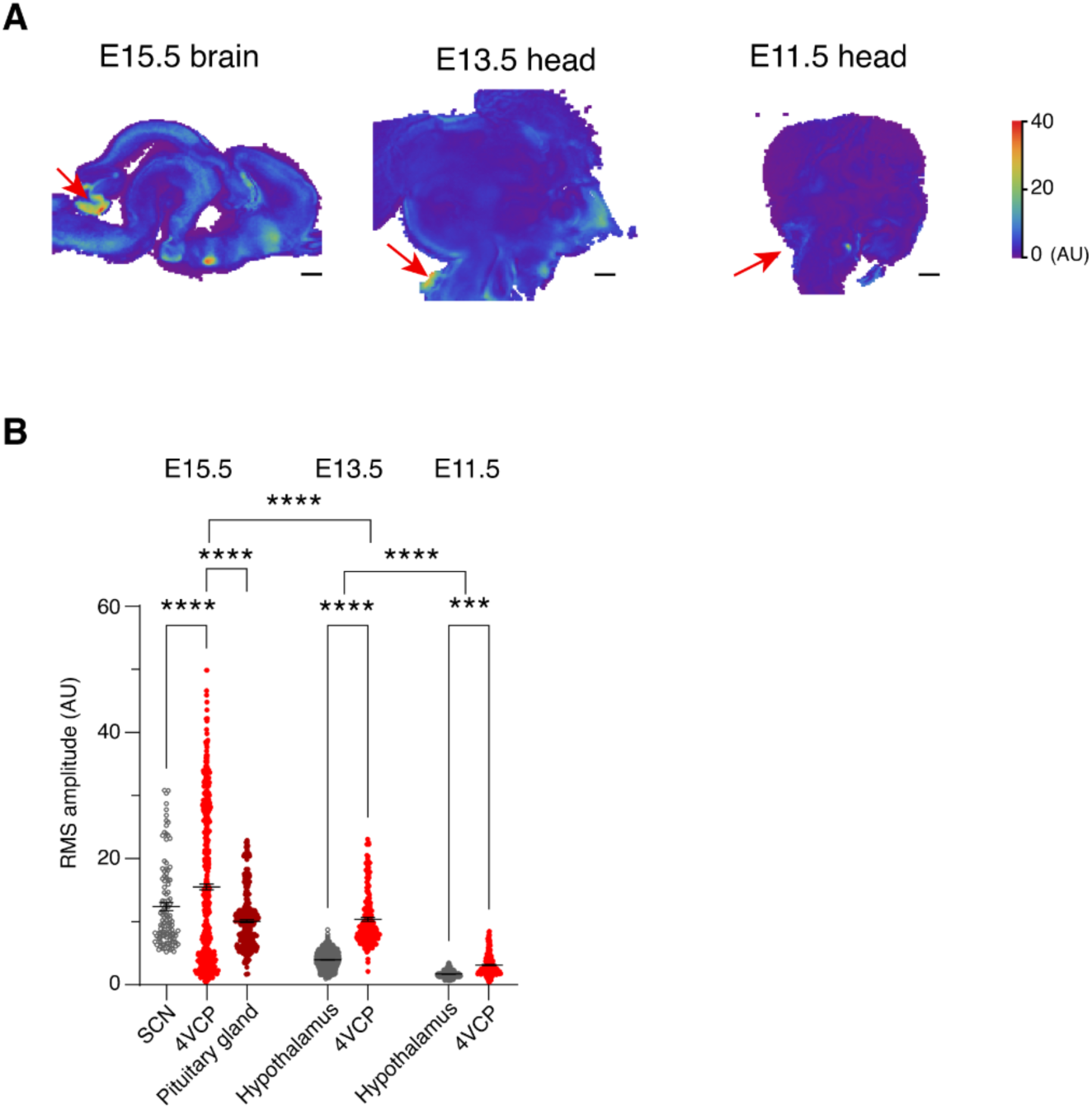
RMS amplitude of PER2::LUC in whole-brain slices at E15.5, E13.5, and E11.5. (**A**) Single-cell level RMS amplitude maps of brain or head slices from E15.5 (left), E13.5 (middle), and E11.5 (right) PER2::LUC embryos. Representative of 3 experiments (5 slices at E15.5, 4 at E13.5, 3 at E11.5). Red arrows indicate the 4VCP. (**B**) RMS amplitude per ROI by brain region within the same slice: SCN (or hypothalamus), 4VCP, and pituitary gland at E15.5, E13.5, and E11.5. One-way ANOVA (*F*(6,3911) = 382.5, *p* < 0.0001) followed by Tukey’s post hoc test (\*\*\*\**p* < 0.0001; \*\*\**p* < 0.001; ns: not significant). Error bars: mean ± SEM from one representative slice per stage, after outlier exclusion. Scale bar: 500 µm.

**Figure S5.**
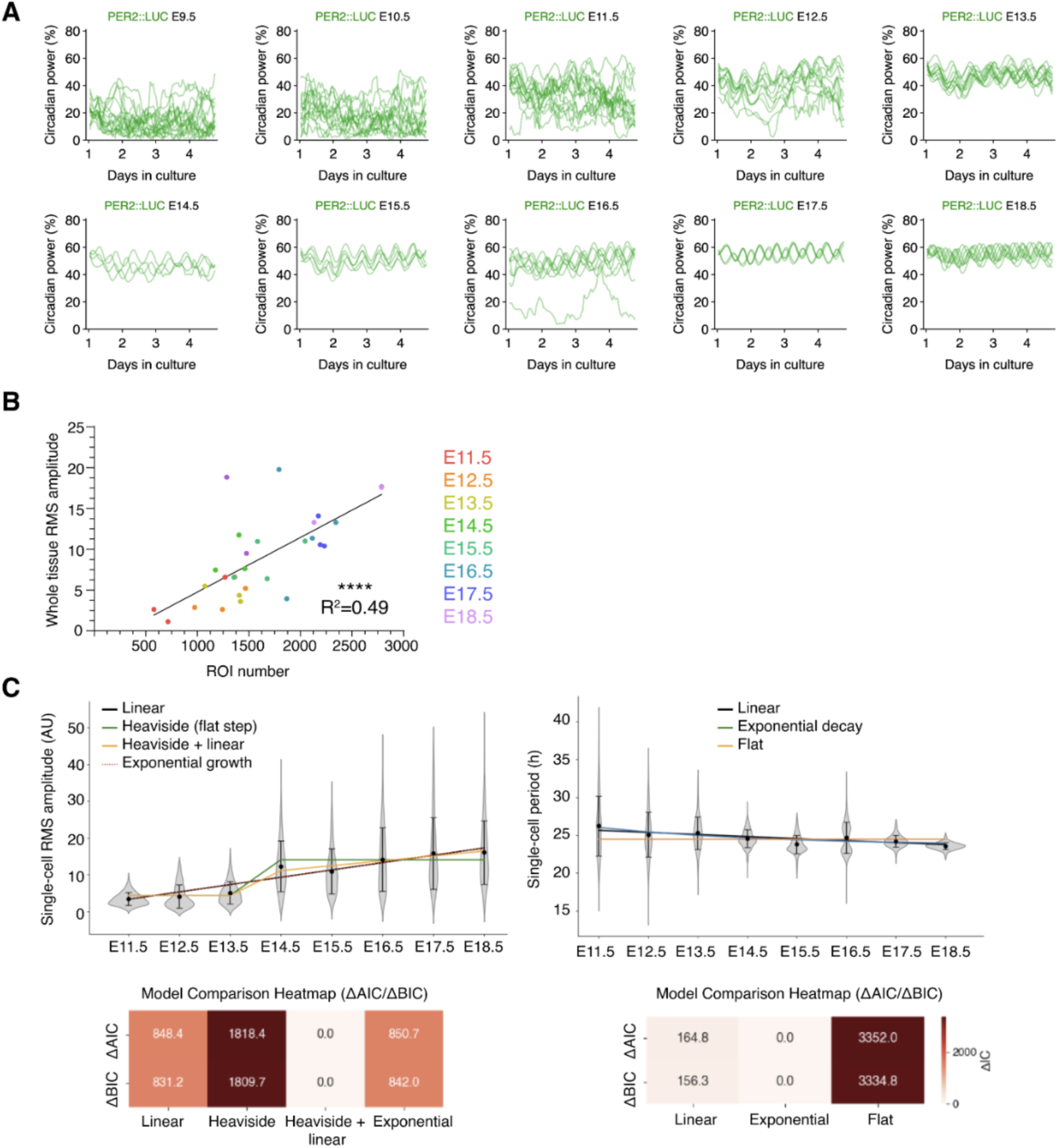
Emergence of significant PER2::LUC circadian power in the 4VCP and relationship between whole-tissue amplitude and cell number. (**A**) Circadian power of individual PER2::LUC 4VCP explants, each line representing one explant. By E12.5, most circadian power trajectories exceed the 20% detection threshold. (**B**) Linear correlation between whole-tissue RMS amplitude and the number of ROIs classified as cells in the embryonic 4VCP across embryonic development (*R*² = 0.49; \*\*\*\**p* < 0.0001). The line shows ordinary least-squares fit. (**C**) Model comparison supporting SNIC-like dynamics in single-cell imaging data. Amplitude (left) and period (right) across stages, fitted with candidate models. Heatmaps (bottom) identify the best-fitting model using Akaike Information Criterion (AIC) and Bayesian Information Criterion (BIC) differences (ΔAIC/ΔBIC) relative to the best model (0 = best; lower = better).

**Figure S6.**
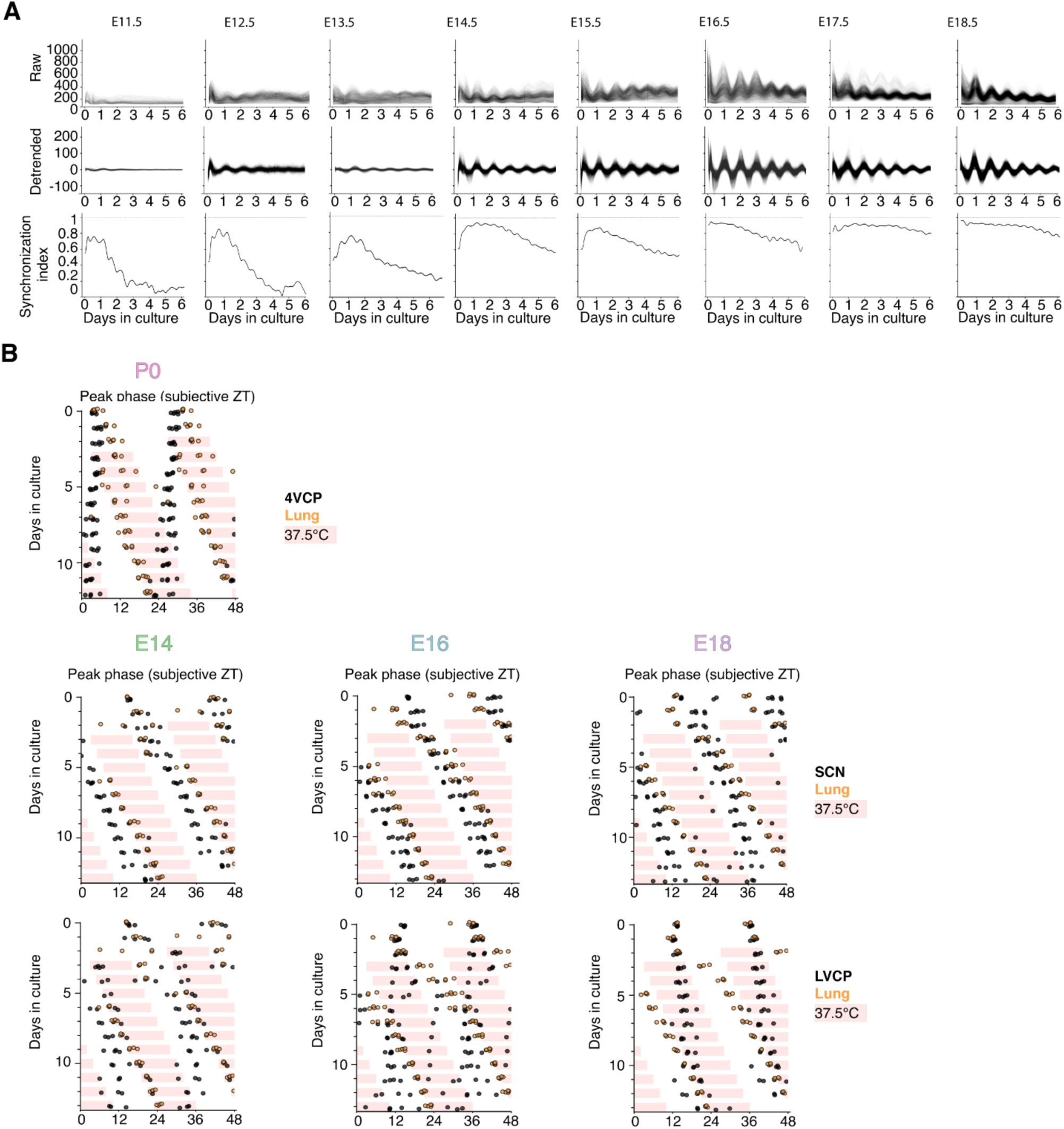
4VCP cellular synchrony across development and sensitivity of the embryonic SCN and LVCP to a 0.5 °C temperature cycle. (**A**) Raw (top) and detrended (middle) PER2::LUC signals from single-cell imaging of the 4VCP from E11.5 to E18.5, with the corresponding Kuramoto order parameter (bottom). Representative of 3-7 experiments. (**B**) Double-plots of PER2::LUC peak phases from neonatal 4VCPs at P0 (*n* = 4-5; top), embryonic SCN (middle), and LVCPs (bottom) at E14, E16, and E18. Explants were recorded for 13 days under a 0.5 °C, 26 h cycle following 2 days at 37.0 °C. Lung explants of the corresponding age are shown in orange as an entrainment reference. Pale pink shading indicates the 37.5 °C temperature epochs.

**Figure S7.**
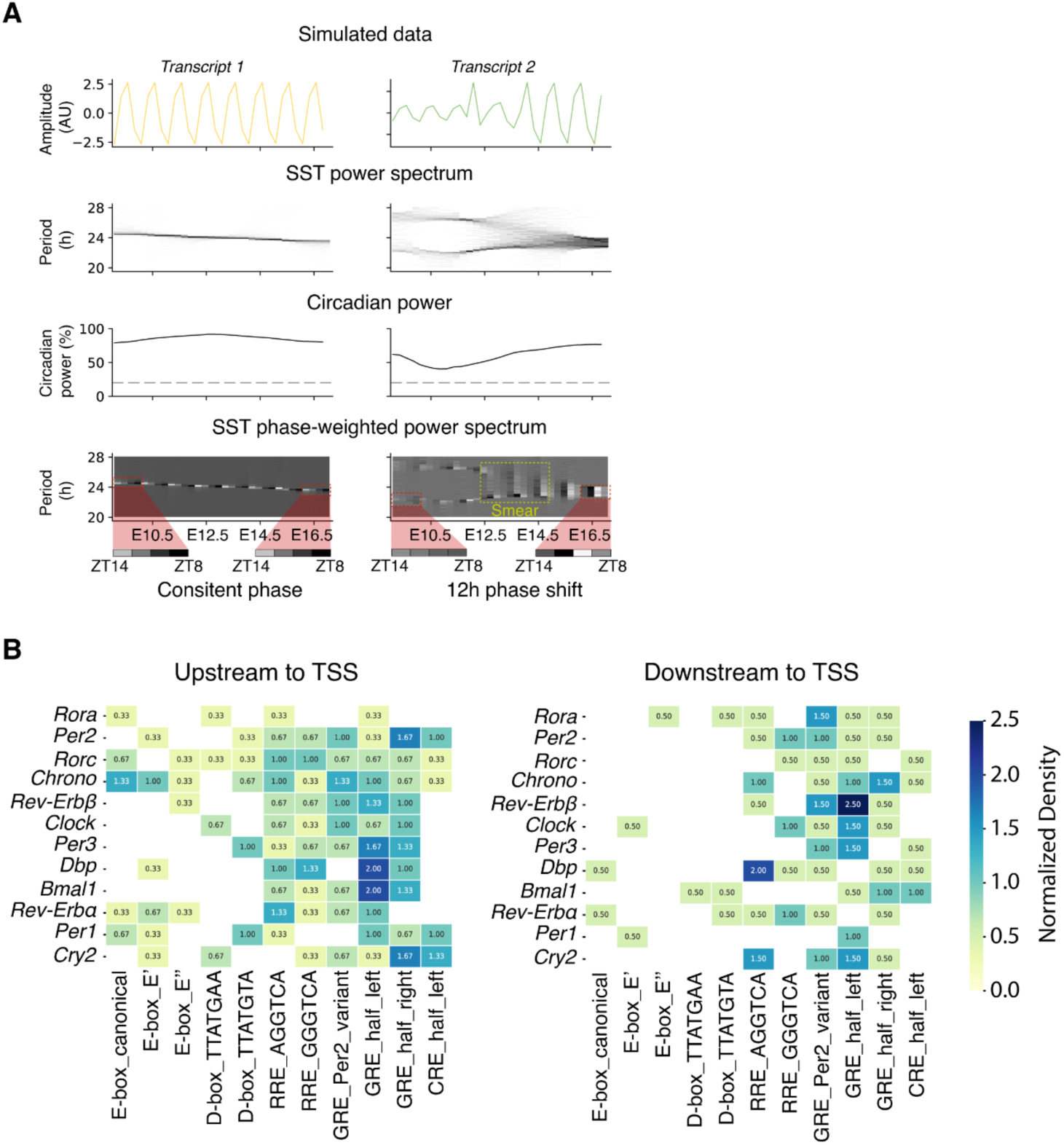
SST-CWT analysis of simulated data and promoter motif density map. **(A)** SST-CWT analysis of simulated time series. Simulated signals for *Transcript 1* (top left) and *Transcript 2* (top right) across developmental time. *Transcript 1* (maternal-like) maintains a stable amplitude and phase; *Transcript 2* (embryonic-like) undergoes amplitude modulation and a 12 h phase shift. SST-CWT power spectrograms highlighting dominant circadian periodicities between 20 and 28 h (upper middle). Circadian power of both simulated transcripts (lower middle). Phase-weighted SST-CWT spectrograms, where color intensity reflects the phase consistency of the dominant period (bottom). Red shading marks early(E9.5) and late (E16.5) windows illustrating phase reorganization; yellow box indicates the spectral smearing interval. **(B)** Normalized density of transcription factor binding motifs for each clock gene.

**Figure S8.**
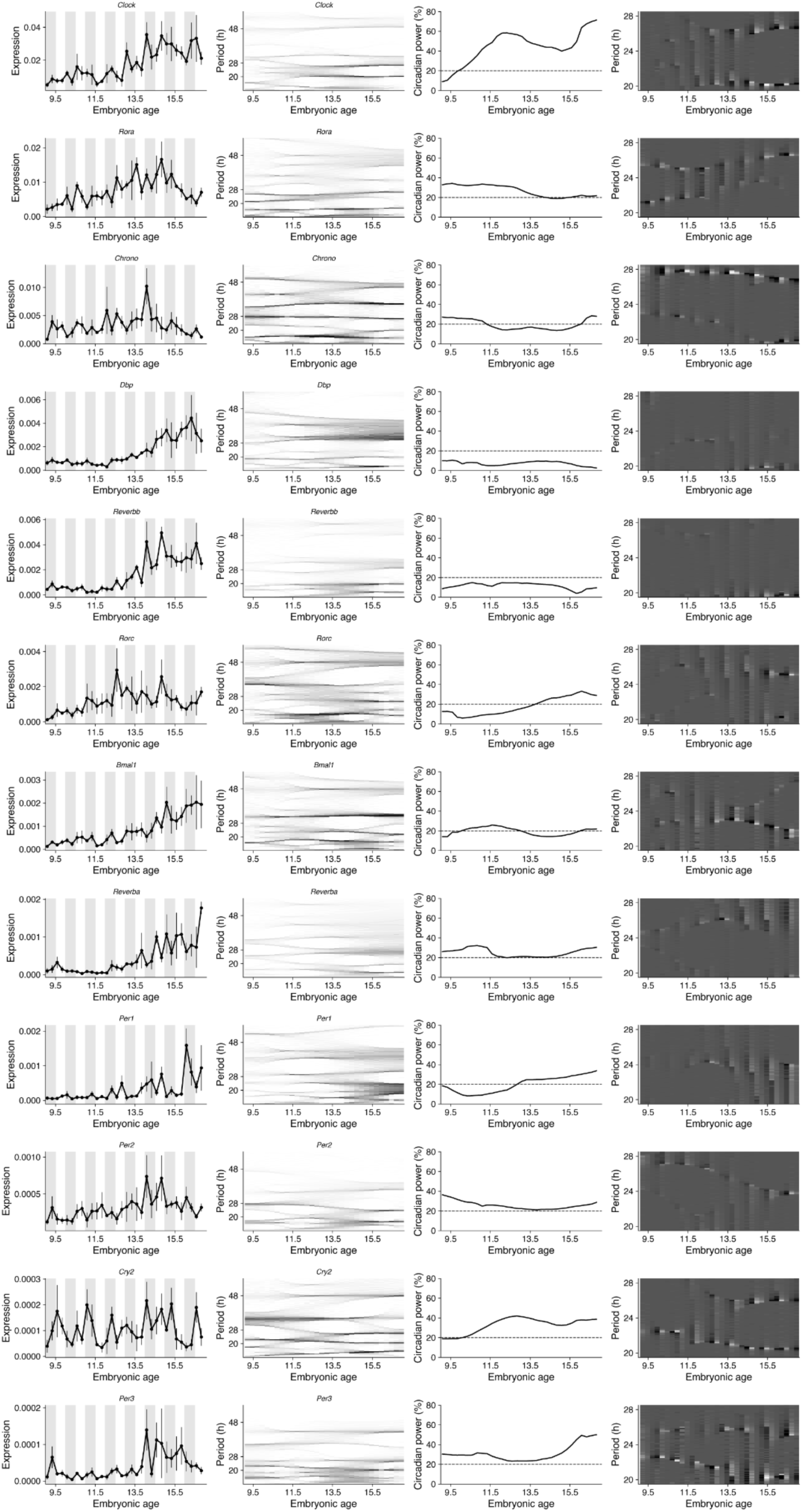
Circadian rhythmicity assessment of embryonic clock genes in the 4VCP. For each gene (rows), four panels are shown from left to right: normalized mRNA expression across embryonic development, sampled every 6 h (ZT2, ZT8, ZT14, ZT20; grey shading indicates the dark phase); SST-CWT spectrogram highlighting dominant circadian periodicities between 20-28 h; circadian power trajectory (dashed line, 20% detection threshold); and phase-weighted SST-CWT spectrogram indicating both dominant period and circadian phase. Error bar: mean ± SEM after outlier exclusion.

**Figure S9.**
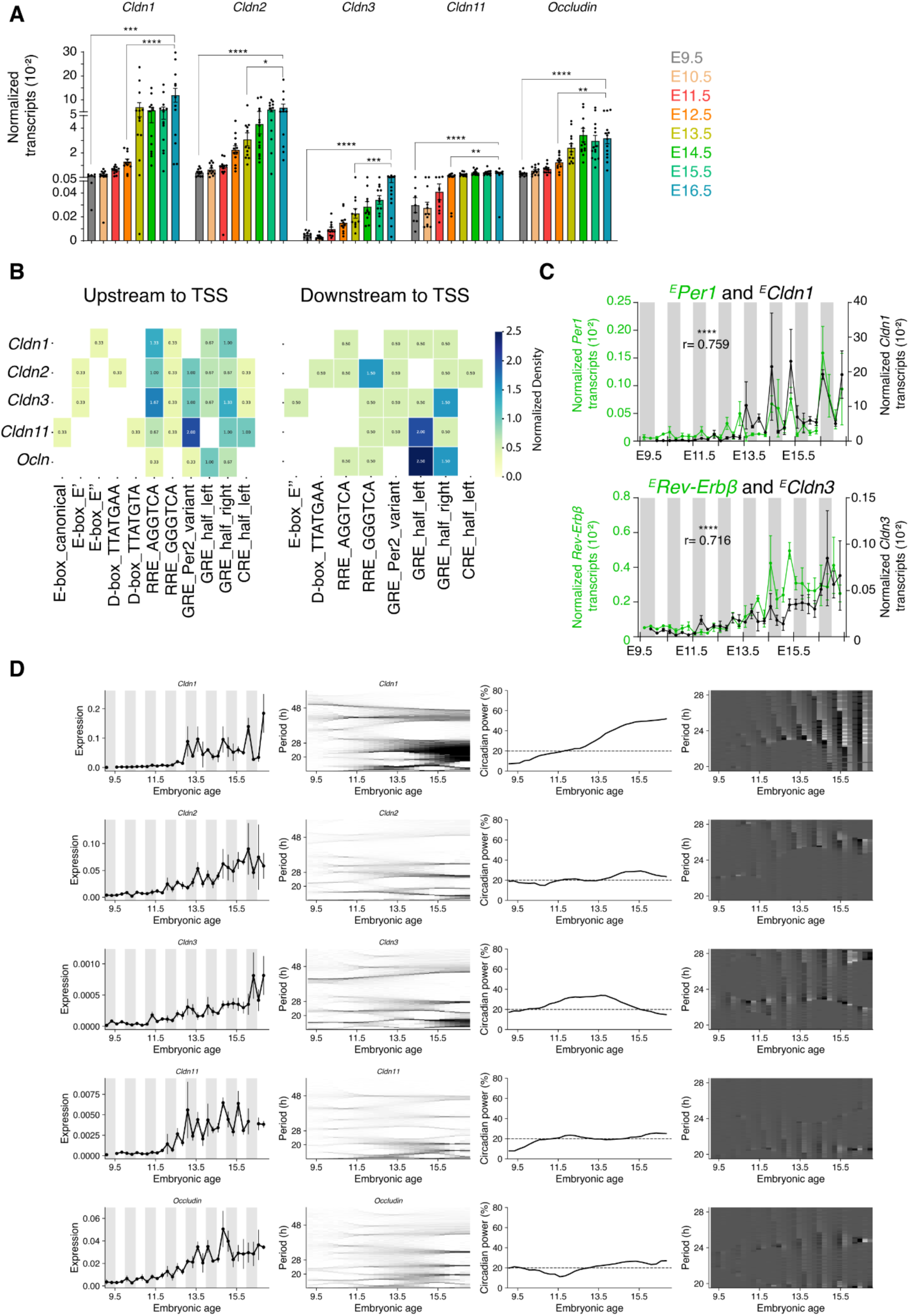
Circadian rhythmicity assessment of embryonic tight junction genes in the embryonic 4VCP. (**A**) Normalized mRNA abundance of tight junction genes in the embryonic 4VCP from E9.5 to E16.5 (*n* = 8-25 litters; 3-11 embryos per litter). One-way ANOVA with Tukey’s post hoc test (\**p* < 0.05, \*\**p* < 0.01, \*\*\**p* < 0.001, \*\*\*\**p* < 0.0001). (**B**) Normalized density of transcription factor binding motifs for each tight junction gene. (**C**) Overlaid mRNA expression of embryonic *Per1* and *Cldn1* (top; *r* = 0.759) and *Rev-Erbβ* and *Cldn3* (bottom; *r* = 0.716) in the 4VCP across development. Pearson correlation; \*\*\*\**p* < 0.0001. (**D**) As in Fig. S8, for tight junction genes. Error bars: mean ± SEM after outlier exclusion.

**Figure S10.**
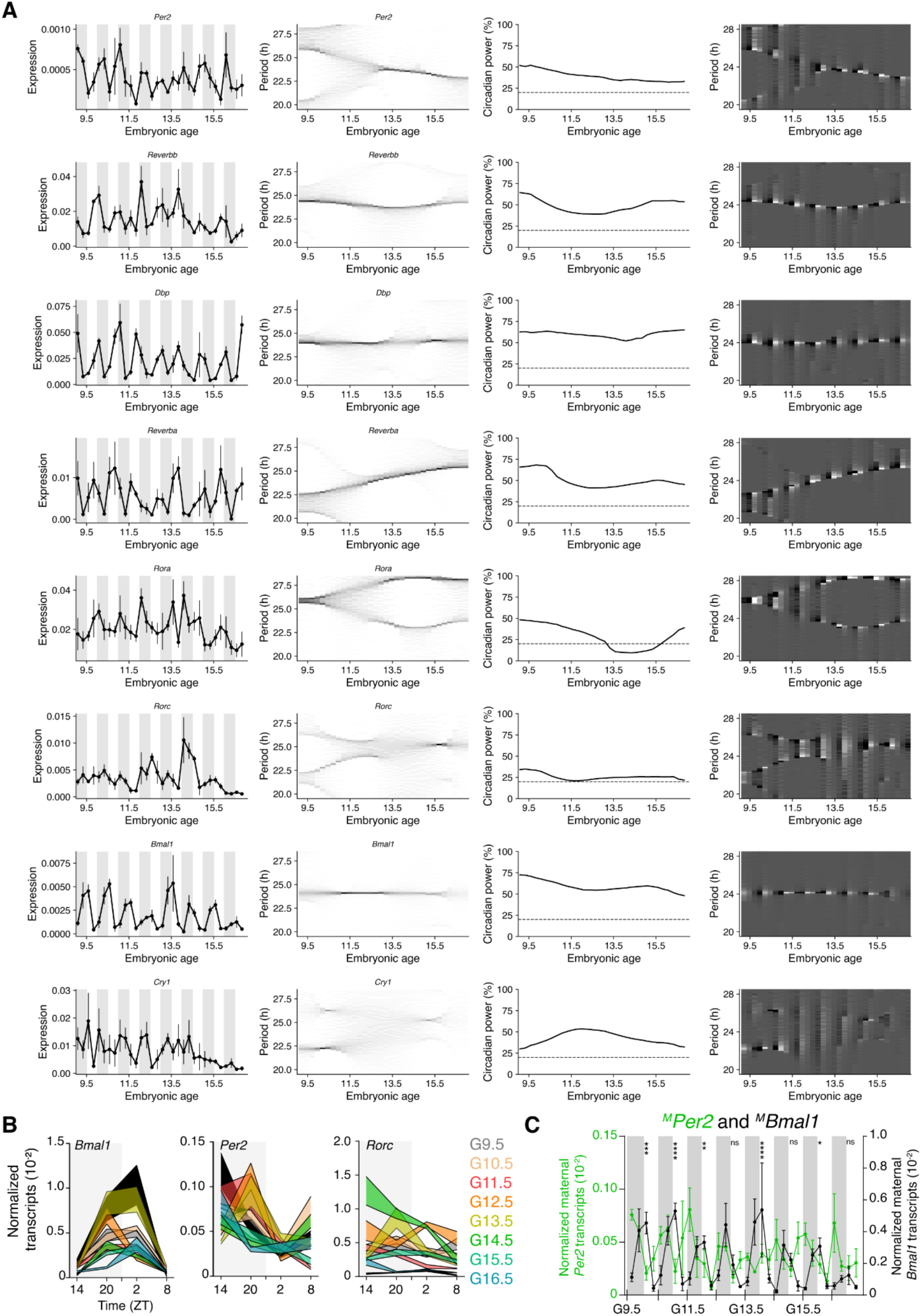
Circadian rhythmicity assessment of maternal clock genes in the 4VCP. (**A**) As in Fig. S8, for maternal clock genes from G9.5 to G16.5. (**B**) Normalized mRNA abundance of maternal *Bmal1*, *Per2*, and *Rorc*, across ZT from G9.5 to G16.5. Curves show mean ± SEM from 2-4 dams or 3-5 non-pregnant females. (**C**) Overlaid normalized mRNA expression of maternal *Per2* (green) and *Bmal1* (black) in the maternal 4VCP across gestation. Two-way ANOVA comparing *Bmal1* and *Per2* expression across gestational days (time effect: *F*(31,131) = 4.072, *p* < 0.0001; gene effect: *F*(1,131) = 135.9, *p* < 0.0001), followed by Šidák’s post hoc test at ZT2 (\**p* < 0.05; \*\**p* < 0.01; \*\*\**p* < 0.001; \*\*\*\**p* < 0.0001; ns, not significant). Error bars: mean ± SEM after outlier exclusion.

**Figure S11.**
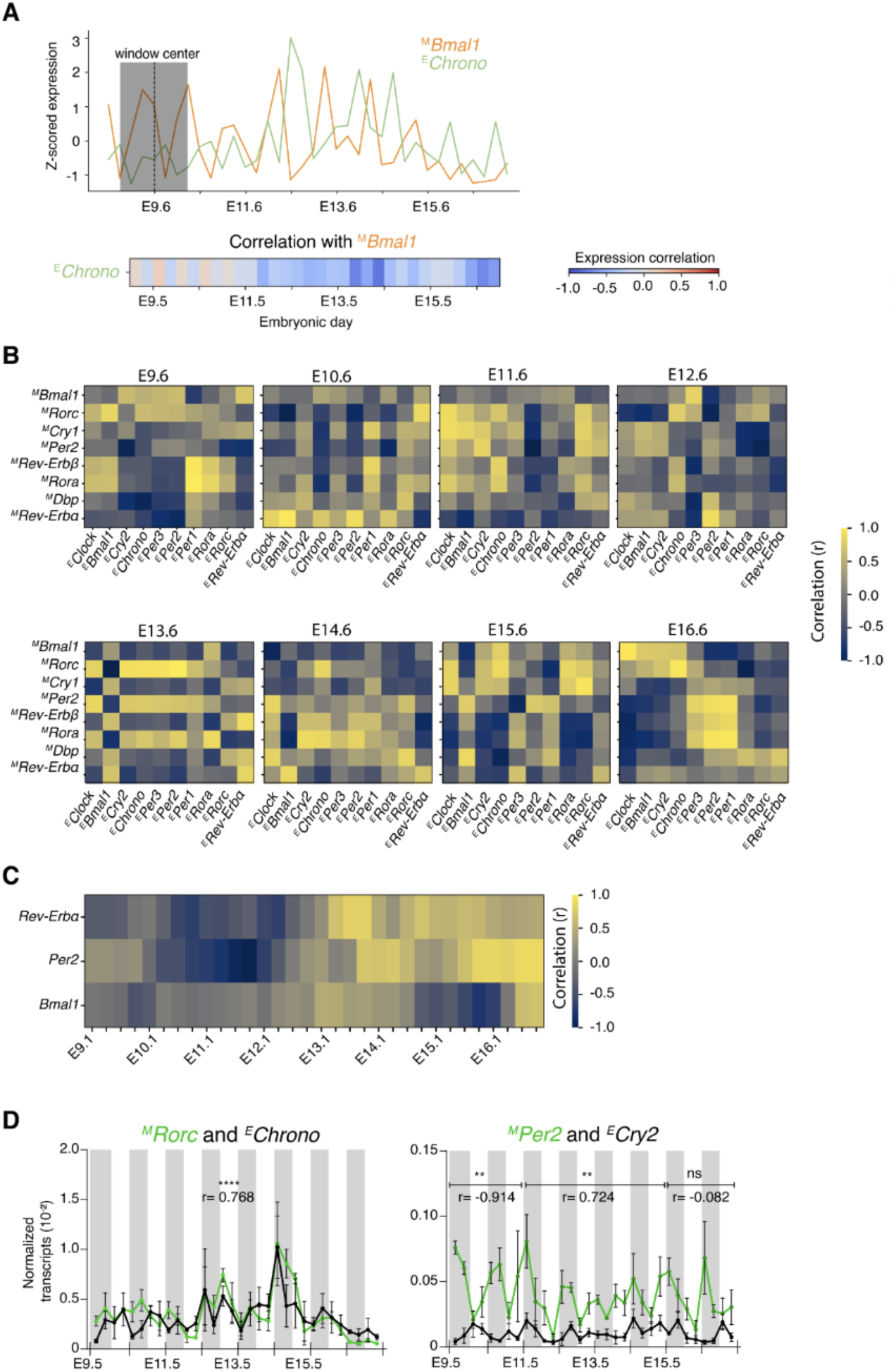
Correlations between maternal and embryonic clock gene expression in the 4VCP. (**A**) Illustration of the sliding-window correlation method between maternal ^M^*Bmal1* and embryonic ^E^*Chrono* expression (top). Resulting correlation trajectory between the two genes in the 4VCP from E9.5 to E16.5 (bottom). (**B**) Pairwise Pearson correlation heatmaps generated using a five-time-point sliding window of transcript levels between maternal and embryonic 4VCP from E9.6 to E16.6. (**C**) Same-gene correlations of *Per2*, *Bmal1*, and *Rev-Erbα* between maternal and embryonic 4VCP from E9.1 to E16.8. (**D**) Overlaid normalized mRNA expression (mean ± SEM) for selected maternal-embryonic pairs: ^M^*Rorc* vs. ^E^*Chrono* (left; *r* = 0.768, \*\*\*\**p* < 0.0001); ^M^*Per2* vs. ^E^*Cry2* (right; negatively correlated before E11.5, *r* = -0.914; positively correlated E11.5-E14.5, *r* = 0.724; not correlated E15.5-E16.5, *r* = -0.082). Pearson correlation; \**p* < 0.05; \*\**p* < 0.01; \*\*\*\**p* < 0.0001; ns, not significant. Error bars: mean ± SEM after outlier exclusion.

**Figure S12.**
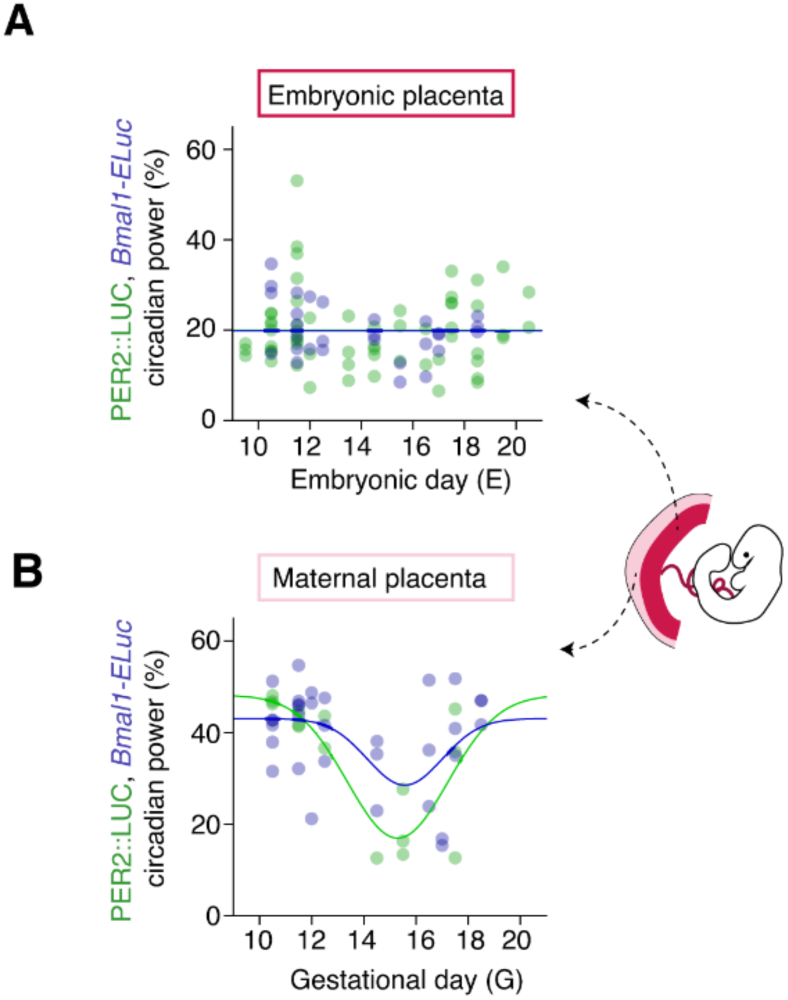
Placental clock robustness during development. (**A**) PER2::LUC and *Bmal1-ELuc* circadian power in the embryonic placenta (dark pink; *n* = 29 and 68, respectively) from E9.5 to E20.5. (**B**) PER2::LUC and *Bmal1-ELuc* circadian power in the maternal placenta (light pink; *n* = 15 and 32, respectively) from G10.5 to G18.5. Lines show inverted Gaussian fits. Schematic (right) illustrates the embryonic (dark pink) and maternal (light pink) placental compartments.

**Figure S13.**
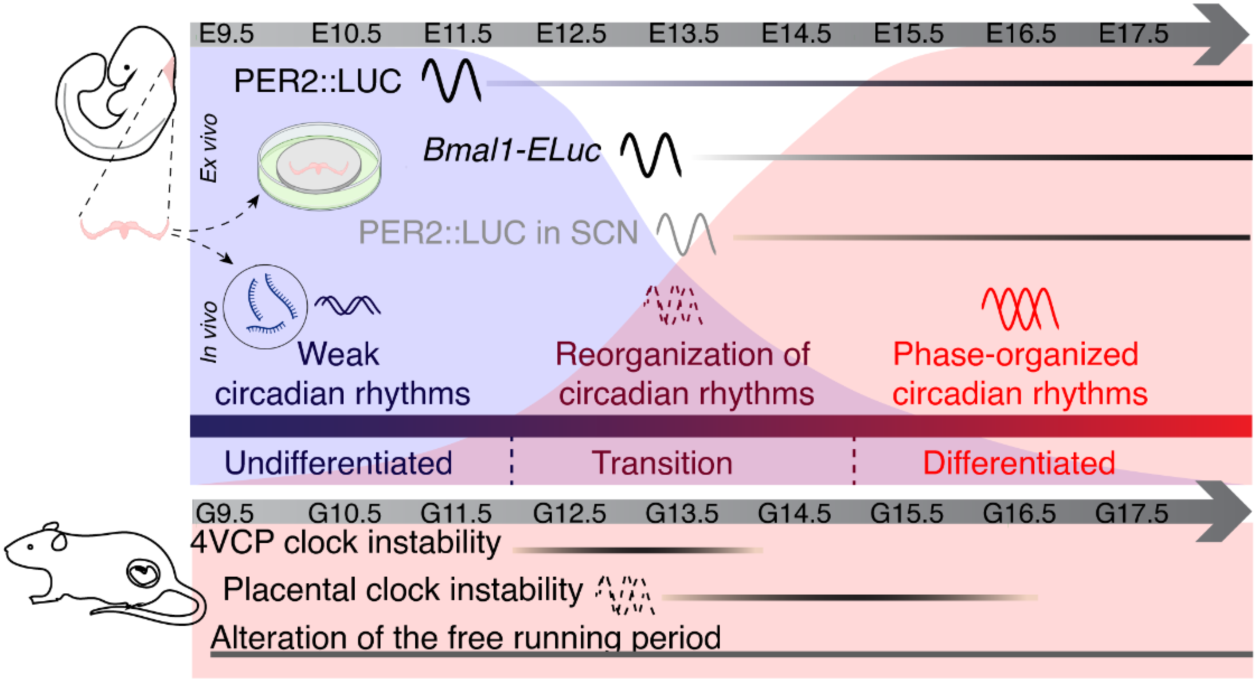
Three differentiation-linked epochs characterize the emergence of the first clock in the embryonic mouse brain. 4VCP is the first brain structure to exhibit autonomous circadian rhythms *ex vivo*, preceding the SCN. Clock assembly is tightly coupled to the tissue’s differentiation program and proceeds through three developmental epochs. **(1) Undifferentiated (E9.5-E12):** Clock genes show weak, non-canonical circadian oscillations *in vivo*, despite the absence of an autonomous clock *ex vivo*. **(2) Transition (E12-E15):** As 4VCP cells differentiate, autonomous PER2::LUC oscillations emerge *ex vivo*, while *in vivo* clock gene rhythms undergo transient destabilization, coinciding with mid-gestational instability in the maternal 4VCP clock, placental clock, and maternal free-running behavior. **(3) Differentiated (>E15):** Robust, phase-organized transcriptional circadian oscillations are established *in vivo*. *Ex vivo*, intercellular synchrony strengthens, and the 4VCP matures into a robust oscillator no longer entrainable by low-amplitude maternal temperature cycles.

## Supplementary Movies

**Movie S1.** Time-lapse of PER2::LUC bioluminescence in an E15.5 sagittal brain slice. Circadian rhythms are detected in both the SCN and the 4VCP. Elapsed time (h) is shown in the upper left. Scale bar: 500 µm.

**Movie S2.** Time-lapse of PER2::LUC bioluminescence in an E13.5 sagittal head slice. A circadian rhythm is detected only in the 4VCP. Elapsed time (h) is shown in the upper left. Scale bar: 500 µm.

**Movie S3.** Time-lapse of PER2::LUC bioluminescence in an E11.5 sagittal head slice. PER2::LUC signal is present in the 4VCP region, but no circadian rhythm is detected. Elapsed time (h) is shown in the upper left. Scale bar: 500 µm.

**Movie S4.** Time-lapse of PER2::LUC bioluminescence in an isolated E13.5 4VCP explant. An autonomous circadian rhythm is evident *ex vivo*. Elapsed time (h) is shown in the upper left. Scale bar: 500 µm.

